# Molecular and phenotypic footprints of climate in native *Arabidopsis thaliana*

**DOI:** 10.64898/2026.03.02.709013

**Authors:** Eneza Yoeli Mjema, Maria Letícia Bonatelli, The NATIVE Consortium, Sascha Laubinger

**Affiliations:** Institute of Biology, Martin Luther University, 06120 Halle (Saale), Germany; German Centre for Integrative Biodiversity Research (iDiv), Halle-Jena-Leipzig, Germany; Institute of Biology and Environmental Sciences, University of Oldenburg, 26129 Oldenburg, Germany; Institute of Ecology, Leuphana Universität, 21335 Lüneburg, Germany; Institute of Agricultural and Nutritional Sciences, Martin Luther University, 06120 Halle (Saale), Germany; Max Planck Institute for Biology, 72076 Tübingen, Germany

## Abstract

Climate change poses a major threat to humanity by driving biodiversity loss and reducing crop yields. To understand the molecular and developmental impacts of rising temperatures, plant science has relied heavily on the model organism *Arabidopsis thaliana*. Despite decades of research, its development under fully natural conditions remains poorly understood, and only approximately 30% of *A. thaliana* genes have experimental functional annotations, probably because many functions are subtle or manifest only in specific laboratory or ecological contexts. Here, we address this gap with a landscape transcriptomic approach that integrates intensive phenotyping and transcriptomic profiling of naturally occurring plants in their native habitats. Across two field sites sampled between 2021 and 2025, we collected >30,000 quantitative trait measurements from more than 3,000 *A. thaliana* plants and generated >1,600 matching transcriptomes. Together with population genomic analyses indicating limited within-population genetic diversity, these data provide an opportunity to investigate phenotypic and transcriptional plasticity associated with natural climatic fluctuations. Year-to-year variation in weather, including temperature anomalies, was associated with differences in plant traits, and climatic variables together explained up to 17% of phenotypic variation. In situ transcriptomes carried clear temperature and local environmental signatures, closely matching temperature-response programs known from the laboratory. Leveraging paired per-plant transcriptomes and phenotypes, we applied machine learning to predict regulators of climate-relevant and other plant traits under natural conditions. The models recovered canonical thermomorphogenesis regulators, including *PHYTOCHROME INTERACTING FACTOR 4* (*PIF4*), providing ecological evidence that temperature signaling pathways defined in controlled environments operate in the wild, and expanded this regulatory landscape by identifying hormonal receptors, signaling components, and previously uncharacterized genes, some of which we functionally validated. Together, this work demonstrates that landscape transcriptomics, by pairing field transcriptomes capturing environmental and regulatory states with phenotypes, enables the predictive identification of genetic regulators of temperature responses and broader plant traits. This makes landscape transcriptomics a scalable framework for climate-aware functional genomics in plants.

## Main Text

Climate change threatens biodiversity and food security by altering environmental conditions that plants must tolerate and adapt to^1–3^. Understanding the molecular mechanisms that enable such adaptive responses requires studying plants in the environments where these stresses naturally occur. The model species *A. thaliana* has yielded unparalleled insights into plant development and stress signaling under controlled conditions, yet its biology under fully natural conditions remains poorly understood. More broadly, recent perspectives have emphasized the need to complement reductionist laboratory approaches with experimental frameworks that interrogate regulatory networks under natural environmental variability^4^. To address this challenge, research has increasingly focused on studying model plant species under more natural conditions^5–10^. The majority of these studies used common-garden experiments, either in controlled conditions or in field-like settings, in which different genotypes were grown under shared environmental conditions^11^. Such settings minimize environmental variation within gardens, allowing trait differences to primarily reflect genetic differences among populations or genotypes. A few field-based common garden experiments have attempted to circumvent this limitation by using local soils and neighboring plants, but only true in situ settings likely reflect authentic real-world trait variation^12,13^. Comparisons between traits measured in field common garden experiments and true in situ conditions remain extremely rare, but some studies suggest that trait variation in situ far exceeds that observed in common gardens^8,14,15^. Furthermore, experimental warming underestimates the extent to which plant traits change in response to rising temperatures^16^. Phenotypic plasticity is often underestimated in common garden experiments, which aim for uniform growth conditions to isolate genetic effects; yet plasticity itself is an adaptive trait under natural selection and is crucial for plants coping with climate variation and ecosystem heterogeneity^17^. Despite these limitations, laboratory experiments and common garden approaches have provided extensive functional data on genes through genome-wide association studies (GWAS) and transcriptome-wide association studies (TWAS) for environmentally influenced traits such as flowering time, seed dormancy, or potential adaptations to future hotter climates^18,19^. In situ phenotyping of *A. thaliana* remains rare, and when available, is typically limited to a few traits and scattered individuals, restricting comprehensive insights into phenotypic plasticity^20,21^. Similarly, field-based transcriptomic studies in *Arabidopsis* and rice have identified genetic background, developmental stage, and environmental conditions including temperature as contributors to transcriptomic variation^22–24^; however, genuine in situ gene expression data remain limited and are available only for a few other plant species.

Although an organism’s transcriptome may not directly reveal gene functions, studying it is nevertheless essential for understanding the molecular strategies that help plants cope with environmental changes, such as those driven by climate change. Across model organisms such as bacteria, yeast, flies, and nematodes, systematic perturbations have enabled functional annotation of most genes^25–28^. In humans, about 80% of genes are supported by experimental functional evidence^29,30^. By contrast, in *A. thaliana*, only about 30% of genes have experimental functional data, while the remainder are inferred by homology or remain uncharacterized^31^. Many of these unknown functions are expected to emerge only under complex or fluctuating environments, which are exactly the conditions plants face in a changing climate. Here, we address this gap with a landscape transcriptomic framework that integrates multi-year field phenotyping with large-scale transcriptomic profiling of *A. thaliana* under natural conditions^32^. We first quantify phenotypic and climatic variation in situ; then we test whether transcriptomes reflect these environments; finally, we leverage paired per-plant transcriptomes and phenotypes to predict candidate regulators of climate-relevant and broader traits. This approach enables the discovery of gene functions that directly mediate plasticity, resistance, and resilience to climatic variability.

### Large-scale field phenotyping reveals key growth principles in Arabidopsis

To better understand natural growth patterns, phenotypic plasticity, and adaptive responses in the model plant *A. thaliana*, we aimed to phenotype a large number of plants that germinated and developed under entirely natural conditions without human intervention at any developmental stage. After identifying suitable natural populations of *A. thaliana*, we conducted extensive in situ phenotyping at two primary study areas: on the East Frisian Island of Spiekeroog, North Sea, Germany, covering about 18.25 km^2^ (53.77°N, 7.73°E), and near the village of Brachwitz in Saxony-Anhalt, Germany, of about 8.34 km^2^ (51.53°N, 11.87°E) (Fig. 1a). Spiekeroog is characterized by a maritime climate with average annual temperatures of 9–10 °C. At Spiekeroog, we sampled each spring from 2021 to 2025 and once in winter 2021. In contrast, Brachwitz experiences a more continental climate with drier summers, where *A. thaliana* was collected during the spring from 2023 to 2025 (Fig. 1c). At both field sites, *A. thaliana* predominantly behaved as a winter-annual species, with most individuals germinating in autumn, completing vegetative growth before winter, and undergoing reproductive growth in spring. Given the predominantly selfing reproductive mode of *A. thaliana*, low levels of heterozygosity and within-site genetic diversity were expected^33,34^. Low-pass whole-genome sequencing of 76 individuals from Spiekeroog and Brachwitz collected in 2021–2023 revealed moderate within-site genetic diversity (π_Spiekeroog_ = 0.17442, π_Brachwitz_ = 0.15983, Fig. S1) and low genetic differentiation between sites (weighted Weir–Cockerham FST = 0.0615). Population clustering analyses (PCA, phylogenetic reconstruction, and ADMIXTURE) identified a large cluster of closely related Spiekeroog individuals, while other Spiekeroog individuals were interspersed with Brachwitz samples or occupied more divergent branches (Fig. 1b, c, Fig. S1-2). Consistent with this, transcriptome-based estimates from 1,053 individuals indicate that populations from Spiekeroog and Brachwitz exhibit low within-site genetic diversity (Fig. S3), suggesting that phenotypic plasticity is likely the main driver of the observed variation at both sites, while acknowledging possible residual genetic structure. In total, 3,442 *A. thaliana* individuals were characterized in situ across spring (2,719 plants) and winter (723 plants) collections. Spring sampling covered both sites across multiple years, enabling comparisons between sites and among years. Winter sampling was conducted once at Spiekeroog, providing an additional within-site seasonal comparison, although without replication across years. We recorded 11 trait-related in situ measurements and seven metadata variables for each plant (Fig. 1d, Data S1). Other traits were calculated as ratios of measured leaf dimensions, such as the ratio of petiole length to total leaf length (petiole length ratio, PLR) and the ratio of leaf length to leaf width (aspect ratio). These ratios describe relative leaf dimensions and facilitate comparisons among naturally growing plants, whose absolute size reflects asynchronous germination and heterogeneous developmental histories as well as genetic and environmental influences. Overall, we analyzed 3,442 individual plants, yielding more than 30,000 trait data points (Data S2).

**Fig. 1:**
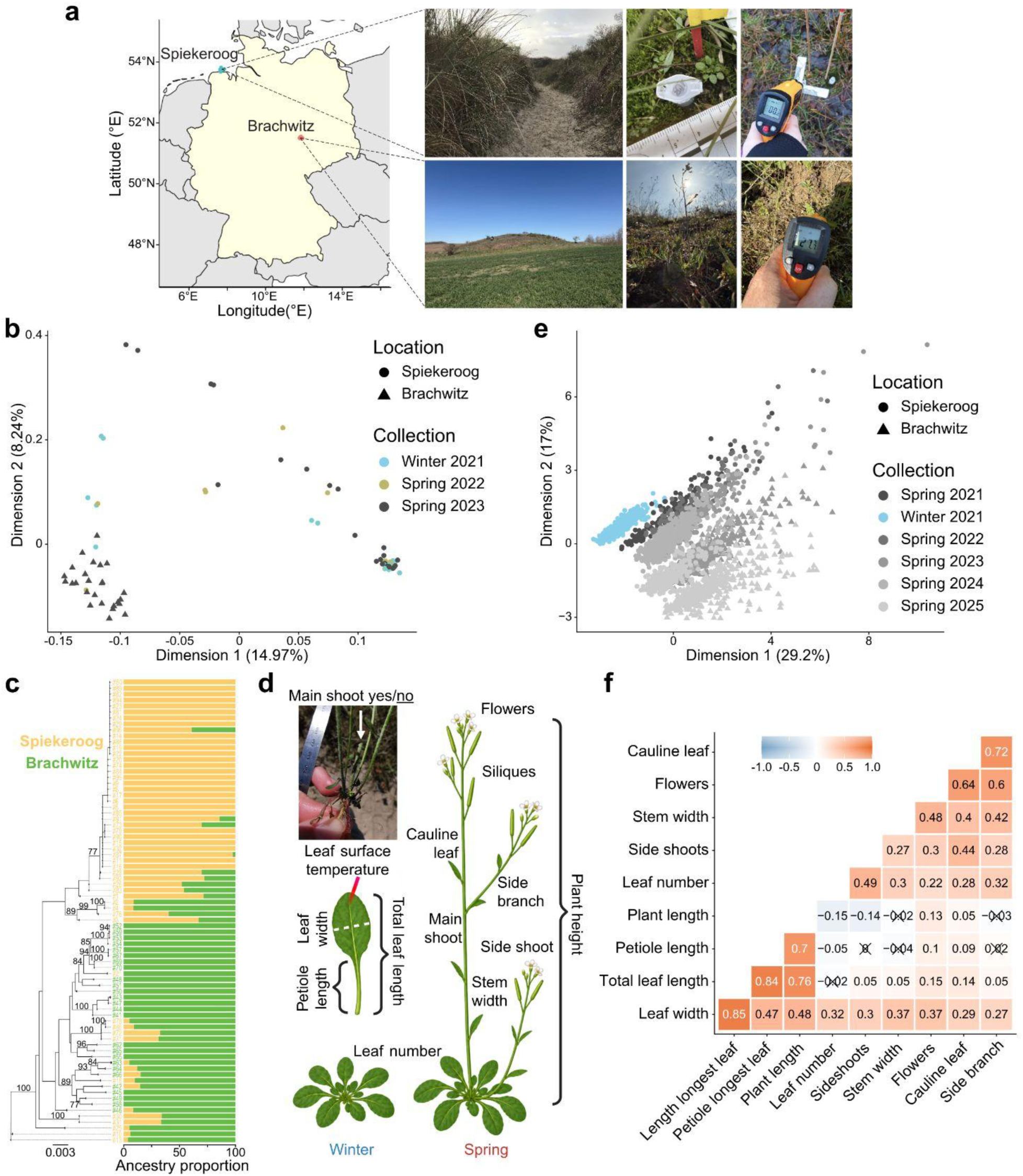
Wild *A. thaliana* displayed seasonal, yearly and location dependent trait variations. **(a)** Geographical map of the study sites in Germany, indicating *A. thaliana* populations on Spiekeroog Island (northern, blue dot) and in Brachwitz (inland, red dot). Adjacent photographs illustrate typical habitats and fieldwork activities, including sampling and temperature measurements. **(b)** A PCA plot showing the genetic clustering pattern of wild *A. thaliana* from Brachwitz (triangular-points) and Spiekeroog (circular-points). Different collections are indicated by different colors as illustrated on the legend. PC1 and PC2 jointly explained 23.21% of the variance with two clusters observed along PC1. **(c)** Genetic structure of the Brachwitz and Spiekeroog accessions. Left: maximum likelihood (ML) tree showing the evolutionary relationships among accessions. Accessions form monophyletic groups by location, with Spiekeroog accessions showing greater divergence from the root. Right: ADMIXTURE bar plots for K = 2 ancestral populations. The ML groups correspond to the two ancestral clusters. Branches with bootstrap values >70 were labeled in the ML tree. Each bar represents one plant. Segment colors indicate inferred ancestry components, and segment lengths indicate their proportions **(d)** Schematic representation of collected morphological traits. During winter, plants were in the vegetative phase (left), and only leaf-related traits were measured. In spring (reproductive phase, right), additional traits such as stem and reproductive structures were recorded. Derived metrics, such as petiole length ratio (PLR) and leaf aspect ratio, were also computed. The presence or absence of a main shoot was documented for all collected plants (top left). **(e)** Biplot of a factor analysis of mixed data (FAMD) based on leaf-related traits, where each point represents a single plant. Colors indicate season–year and shapes indicate site. Dimensions 1 and 2 explain 29.2% and 17.0%, respectively (46.2% combined). Plants cluster primarily by season–year, with winter 2021 (sky blue) forming a distinct group. Variation in leaf traits is further reflected in the spread of individuals within each collection. **(f)** Pairwise Spearman’s correlation of estimates from trait-trait mixed models accounting for the random effect of collection year and location for all spring samples. The heatmap displays both significant (p < 0.05) and non-significant (crossed) correlations between all measured traits. Strong positive correlations were observed among leaf-related traits (r = 0.7–0.85), mirroring the patterns in both winter and spring collections from Spiekeroog and Brachwitz for each year (Fig. S9-S10).

To investigate how season, location, and collection year influence the overall clustering of phenotypic traits, we applied several dimensionality reduction methods to leaf-related traits measurable in both winter and spring. Since classical Principal Component Analysis (PCA) cannot directly incorporate categorical variables, we performed PCA separately for subsets based on these categories (Fig. S4). In contrast, Factor Analysis of Mixed Data (FAMD), a method integrating both quantitative traits and categorical variables (season, location, and year), revealed distinct clusters; this suggests that season, location and year contributed to the separation in our collection (Fig. 1e). As expected, the clear separation along Dimension 1 indicated that both the collection year and season (winter vs. spring) made significant contributions as primary factors driving phenotypic variation (Fig. S5). Motivated by the strong seasonal signal in the clustering, we directly compared leaf-related traits of plants phenotyped in winter 2021 with corresponding plants from the following spring 2022. The number of leaves produced until December (mean = 9.674 leaves, n = 684) differed only moderately, but not significantly (*p* = 0.141), between seasons and increased in spring (mean = 10.84 leaves, n = 125) (Fig. S6d). This suggests that much of leaf initiation precedes winter, whereas elongation dominates in spring. While this notion holds for leaf and petiole length, for which we observed a significant increase from winter to spring (*p* < 2.00e-10), leaf width did not significantly change (*p* = 0.064), suggesting that leaf growth during spring mainly occurs longitudinally rather than laterally (Fig. S6a-c). Extending our analyses beyond leaf morphology to architectural traits in spring collections, Brachwitz and Spiekeroog show significant yearly variation in these traits (Fig. S7 and S8). Taken together, our results suggest yearly differences in vegetative and reproductive growth dynamics among wild *A. thaliana*.

Next, we focused our analysis specifically on spring samples, as these allowed us to evaluate not only vegetative traits but also traits associated with the reproductive growth stage. To identify site-specific trait relationships, we performed pairwise correlation analyses between ten phenotypic traits across eight distinct groups (site-year combinations): Spiekeroog (2021, 2022, 2023, 2024, 2025) and Brachwitz (2023, 2024, 2025). Rosette leaf traits generally showed strong, significant correlations (r > 0.5) within spring samples, similar to those observed in winter samples (Fig. S9 and S10); however, we noted some variation in trait correlations specific to plants in reproductive stages across different years and locations. Notably, the number of leaves did not significantly correlate with plant height across all groups in most years. In contrast, flower number consistently correlated positively with plant height in seven of the eight groups (r = 0.35-0.71, Fig. S9 and S10), and stem width correlated positively with plant height across six groups (r = 0.36-0.59). To confirm whether the observed spring yearly-site correlation persisted after accounting for the random effects of year and location, we calculated pairwise correlations from the trait-trait mixed models. Strong positive correlations (r = 0.7-0.85) among leaf-related traits highlighted coordinated growth in spring similar to patterns observed in winter (Fig. 1f). Interestingly, the number of flowers, cauline leaves, side branches and stem width showed significant positive correlations (r = 0.4-0.7), reflecting a consistent coupling among architectural traits.

We also observed that approximately 7% of all plants lacked a main shoot, most likely due to herbivory^35^ (Fig. 1d, Fig. S11a). Plants without a main shoot produced significantly more side shoots (Fig. S11b), consistent with the classical phenomenon of apical dominance, in which auxin produced in the main shoot suppresses the outgrowth of lateral shoots. No significant difference was observed in flower number (Fig. S11c); however, plants with a main shoot had significantly more siliques, reflecting their developmental head start and the additional time available to produce mature fruits (Fig. S11d). Consequently, plants with a main shoot achieved slightly higher overall reproductive output (as measured by flowers + siliques, Fig. S11e). Importantly, plants that lost their main shoot were nevertheless able to restore reproductive output to ∼83% of that of intact plants, demonstrating that the release of apical dominance functions as a fail-safe against herbivory and can partially compensate for tissue loss. This illustrates that in situ trait variation also reflects biotic perturbations and that compensatory plasticity can buffer fitness under natural damage.

### Temperature anomalies reshape trait variation in the field

Laboratory and common garden experiments with *A. thaliana* provide insights into the effects of climate on development, but in the field, trait variation also arises from genetic, edaphic, biotic, and stochastic factors that cannot be systematically controlled. Nevertheless, our field sites exhibited marked interannual climatic variation in different variables such as soil and air temperatures, precipitation, relative humidity, and sunshine duration, as recorded by nearby weather stations for both locations (Data S3, Fig. S12 and S13). Mean monthly temperatures in February and March fluctuated by up to ∼5 °C between years, with February-May 2024 being particularly warm (anomalies > +2 °C) and the same period in 2021 unusually cold (anomalies < −2 °C) (Fig. 2a, Data S4, Fig. S14 and S15). We next asked whether these climatic anomalies were mirrored in our phenotypic data. Consistent with the well-established effect of high temperatures on petiole elongation^36–38^, plants grown in warmer 2024 had significantly longer PLR than plants grown in colder 2021 (Fig. 2b, Fig. S16) and an overall positive correlation of field petiole length measurement and ambient air temperature across all years and collection sites (Spearman’s r = 0.69, adjusted P = 0.0694, Fig. S17), indicating that interannual temperature anomalies translate into measurable trait differences.

**Fig. 2:**
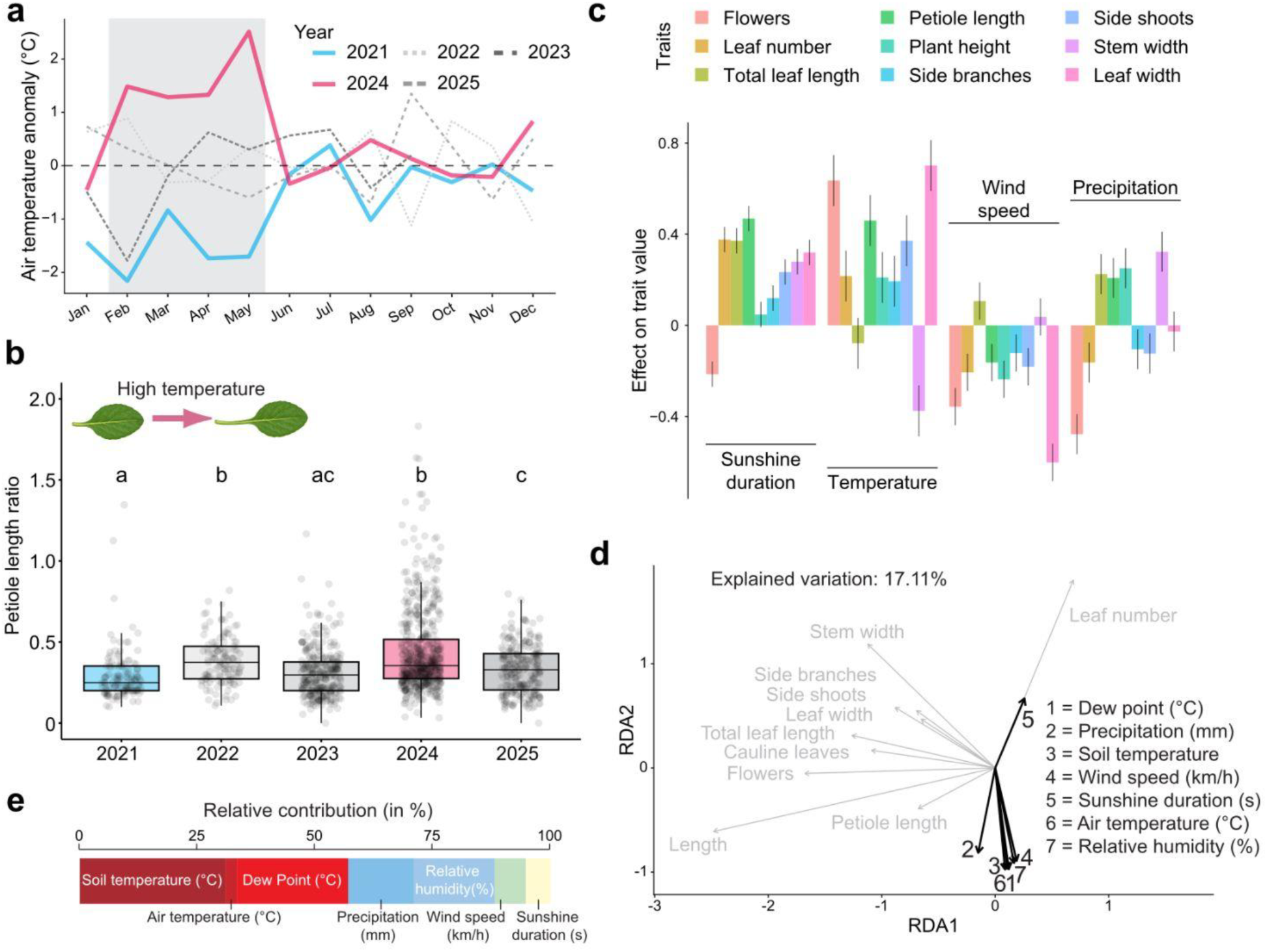
Annual climatic variation contributes to phenotypic diversity in wild *A. thaliana* populations. **(a)** Interannual variation in air temperature at Spiekeroog from 2021 to 2025. Monthly temperature anomalies were calculated as deviations from the multi-year monthly mean (e.g., 2021 to 2025 for Spiekeroog). Pronounced interannual fluctuations were observed, reaching up to ±2 °C in February and March of 2021 and 2024. **(b)** Elevated air temperatures are associated with increased leaf elongation in natural *A. thaliana* populations. Box plots show the petiole length ratio (PLR) of plants collected at Spiekeroog between 2021 and 2025. Plants grown during the warm spring of 2024 exhibited significantly higher PLR values (p < 0.0001) than those from 2021, which experienced the coldest spring temperatures (February–May). **(c)** Morphological traits interacted differently with weather variables. A bar plot of weather-trait mixed-model estimates showing the strength and direction of the weather-trait relation. Trait values were scaled to mean zero and unit variance prior to modeling. Petiole length (green bar) responds positively to air temperature, precipitation, and sunshine duration; conversely, wind speed was negatively associated with most trait estimates. **(d)** Relationship between climatic factors and morphological traits inferred from redundancy analysis (RDA) of all spring samples. The biplot displays associations between trait variables (grey) and climatic predictors (black). Climate parameters collectively explained 17.1% of the total trait variation. Arrow orientation indicates correlation between predictors and traits, with arrow lengths reflecting the relative contribution to the RDA axes. **(e)** Relative contribution of climatic factors to annual trait variability in wild *A. thaliana* populations. Bar plot showing percentage contributions of the seven climatic variables included in the RDA. Temperature-related variables (soil, air, and dew point) accounted for the largest share (57%), followed by precipitation (14%) and relative humidity (17%).

To systematically test trait-weather relationships, we fitted linear mixed models quantifying the influence of each weather variable on individual trait variation across collection years and sites, thereby disentangling trait-specific sensitivities. Petiole length showed strong positive associations with air temperature, sunshine duration, and precipitation, whereas air temperature negatively affected stem width. Wind speed exerted a significantly negative effect on most traits, with the exception of stem width, which was positively influenced by wind speed (Fig. 2c; Data S5).

Motivated by these results, we asked to what degree weather variables explain overall trait variation in the full dataset. Leveraging our multi-year, two-site design, we applied redundancy analysis (RDA) to partition the contribution of climatic factors. Climate explained ∼17% of trait variation, notably given uncontrolled microclimate, edaphic, and biotic variation (Fig. 2d). This indicates that climatic effects are substantial yet heterogeneous, with trait-specific responses to varying climates that are only partially coordinated across traits (Fig. 2d). Among the predictors, temperature-related variables contributed the largest share (>50%), with precipitation (13.9%) and relative humidity (17.2%) also explaining substantial fractions (Fig. 2e; Data S6). Together, these findings demonstrate that, under fully natural conditions, climatic fluctuations directly shape *A. thaliana* phenotypes, providing ecologically relevant insight into plant responses to ongoing climate change.

### Natural transcriptomes encode signatures of temperature and local environment

Because temperature-related weather variables (air, soil, dew point temperatures and sunshine duration) were the dominant climatic drivers of phenotypic variation in our dataset, we next asked whether transcriptome profiles also captured these environmental signatures. As a first step, we analyzed leaf transcriptomes across distinct seasonal environments and developmental phases to test whether natural expression profiles reflect ambient environmental regimes. In total, we generated 1,611 field transcriptomes, including 704 plants collected in winter 2021 at Spiekeroog, and 679 and 228 plants from spring 2022, 2023, and 2024 at Spiekeroog and Brachwitz, respectively. Field sampling was largely restricted to daytime hours (8:30 am – 17:30 pm); accordingly, diurnally regulated genes may contribute to within-day expression differences, and transcripts peaking at night or early morning may be underrepresented. Most sequencing reads could be mapped successfully to the *A. thaliana* genome; however, 10-30% of reads were unmappable, with higher proportions in spring than in winter (Fig. 3a, Fig. S18). Closer inspection revealed that many of these unmapped reads originated from diverse fungal and oomycete species colonizing leaves as endophytes or epiphytes (Fig. 3b). *Albugo*, a genus of plant-parasitic oomycetes, accounted for many of the taxonomically assigned reads (Fig. 3b)^21^. The low abundance of unmapped reads in winter suggests that many of these taxa overwinter as spores or in nearby plants and proliferate in spring, indicating a seasonal restructuring of the leaf-associated microbiome.

**Fig. 3:**
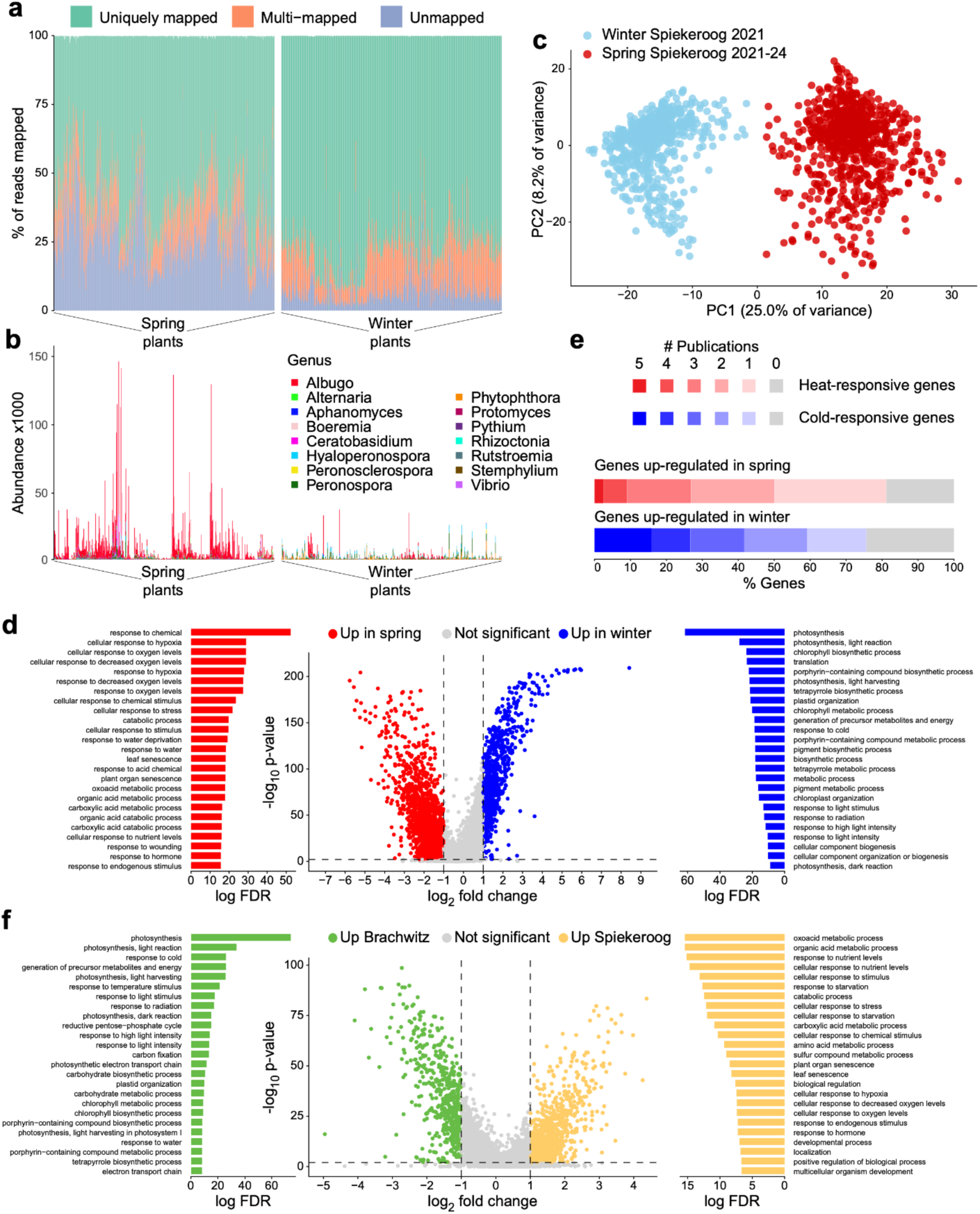
Seasonal and locational transcriptomic signatures in natural *Arabidopsis thaliana* populations. **(a)** Stacked bar plot showing the proportion of reads uniquely mapped to the *Arabidopsis thaliana* reference genome, multi-mapped reads, and unmapped reads for individual samples. On average, ∼8% of reads were unmapped in winter samples and ∼24–26% in spring samples. **(b)** Microbial community composition accessed through the taxonomical classification of reads unmapped to the *A. thaliana* reference genome. **(c)** Distinct transcriptomic profiles separate winter and spring plants. Principal component analysis (PCA) of normalized, batch-corrected gene expression data from Spiekeroog. PC1 (25%) and PC2 (8.2%) explain most of the observed variance. **(d, f)** Differentially expressed genes (DEGs) reveal seasonal and site-specific transcriptomic patterns. Volcano plots showing significantly regulated genes (FDR < 0.01 and |log2FC| > 1). Comparison between winter and spring plants highlights enrichment of processes related to cold and light response in winter, and to senescence, wounding, and abscisic acid (ABA) signaling in spring (c). Comparison between sites reveals enrichment of genes related to hypoxia and nutrient response in Spiekeroog, and to high-light and cold responses in Brachwitz (d). **(e)** Seasonal differences in gene expression closely matched those from temperature shifts in controlled laboratory settings. A stacked bar chart showing the proportion of up-regulated genes in winter (blue) and spring (red) that were reported in at least one of five cold-treatment or heat-treatment studies, respectively. Approximately 75-80% of the genes that were significantly differentially regulated by the seasonal shift were reported in at least one publication in our comparative analysis.

We detected >30,000 expressed genes across all plants, with ∼10,000 consistently expressed in every sample. PCA revealed a clear separation between winter and spring samples along PC1, which explained 25% of the total variance (Fig. 3c). To test whether this divergence reflected differences in temperature rather than developmental stage, we performed a differential expression analysis between winter and spring plants from Spiekeroog. We identified 3,075 significantly differentially expressed genes (DEGs; FDR < 0.01 and |Log_2_FC| > 1), of which 897 were upregulated and 2,178 downregulated in winter (Fig. 3d, Data S7). Winter-upregulated genes were enriched for cold acclimation, low-light response, and photosynthetic light-harvesting processes, consistent with an acclimation strategy to maximize light use under cold, low-radiation conditions (Fig. 3d, Data S8). In contrast, genes associated with abiotic stress and heat shock factors (HSFs) were strongly expressed in spring (Fig. 3d, Data S8). Notably, approximately 75-80% of seasonally regulated genes also responded to thermal shifts in laboratory experiments, supporting the ecological validity of our in situ observations (Fig. 3e). Alongside temperature responses, seasonal expression differences may reflect developmental state and the contrasting sampling strategies: winter samples comprised whole aerial material, predominantly young leaves but potentially including meristematic tissue, whereas spring samples were restricted to the fully developed largest leaf. Consistent with a developmental contribution, winter-upregulated genes included the juvenility-promoting factor *TOE1*, whereas spring-enriched genes included *SPL4*, *FT*, *FTIP1*, FUL and the master senescence regulator *ORE1*, without a corresponding differential-expression signature of canonical shoot stem-cell regulators such as *WUS* and *CLV1–3* (Data S7).

We also examined whether additional local factors shape gene expression. Comparing spring transcriptomes from Spiekeroog (n = 675) and Brachwitz (n = 221) revealed 1,659 significant DEGs (FDR < 0.01 and |Log_2_FC| > 1) between the two sites (Fig. 3f, Data S9). Of these, 1,144 were upregulated in Spiekeroog and 515 in Brachwitz, suggesting that, despite genetic differences, distinct transcriptomic programs exist between the coastal and inland environments. Genes upregulated in Spiekeroog were enriched for hypoxia and senescence-related processes, consistent with potential stress generated by high soil moisture and salinity, whereas Brachwitz samples showed enrichment for light- and radiation-responsive pathways, reflecting growth in more exposed inland habitats (Fig. 3f, Data S9). Taken together, these results confirm temperature as the principal determinant of seasonal transcriptomic variation, while also revealing additional local environmental footprints on gene expression under fully natural conditions.

### Field transcriptomes enable the prediction of candidate genes associated with climate-relevant traits

Climate change is altering temperature regimes worldwide, and plant survival depends on the capacity to sense and respond to such fluctuations. Our data show that *A. thaliana* transcriptomes in the wild carry clear temperature signatures, indicating that field expression profiles reflect the plant’s immediate thermal environment. Whereas the seasonal differential expression contrasts above capture broad environmental and developmental differences (winter vs. spring), they cannot resolve the fine-scale transcriptional variation experienced by individual plants. To overcome this limitation, we exploited per-plant transcriptomes paired with per-plant trait measurements. This design allows us to harness the subtle expression differences arising from each plant’s local temperature exposure and microenvironment, and to link these individualized expression profiles directly to climate-relevant traits such as leaf surface temperature or petiole length. Because transcriptomes reflect a plant’s current regulatory state, their predictive power is expected to be strongest for traits that mirror immediate physiological or environmental conditions. Leaf surface temperature fulfills this criterion: although influenced by longer-term architectural adjustments such as petiole elongation and hyponasty, it is strongly determined by ambient temperature, radiation, and transpiration at the time of sampling. In contrast, traits such as petiole length integrate developmental responses over extended time windows and may therefore be less tightly coupled to the instantaneous transcriptomic state of a single leaf. Traits regulated in other tissues, such as floral traits, are likewise expected to show weaker coupling to leaf transcriptomic variation. Nevertheless, previous work has demonstrated that leaf expression profiles can correlate with later phenotypic outcomes^39,40^.

To systematically test trait–transcriptome predictive relationships under natural conditions, we implemented a machine learning framework combining gene selection, model training, and independent validation (Fig. 4a). In contrast to differential expression analyses, which prioritize transcripts showing condition-dependent changes in abundance, this predictive framework does not require candidate genes to be transcriptionally responsive themselves. Instead, it identifies genes whose expression covaries with trait variation, including upstream regulatory components (e.g., transcription factors, RNA-binding proteins, epigenetic modifiers, and signaling components) and downstream responsive or effector genes (e.g., heat shock proteins or metabolic enzymes).

**Fig. 4:**
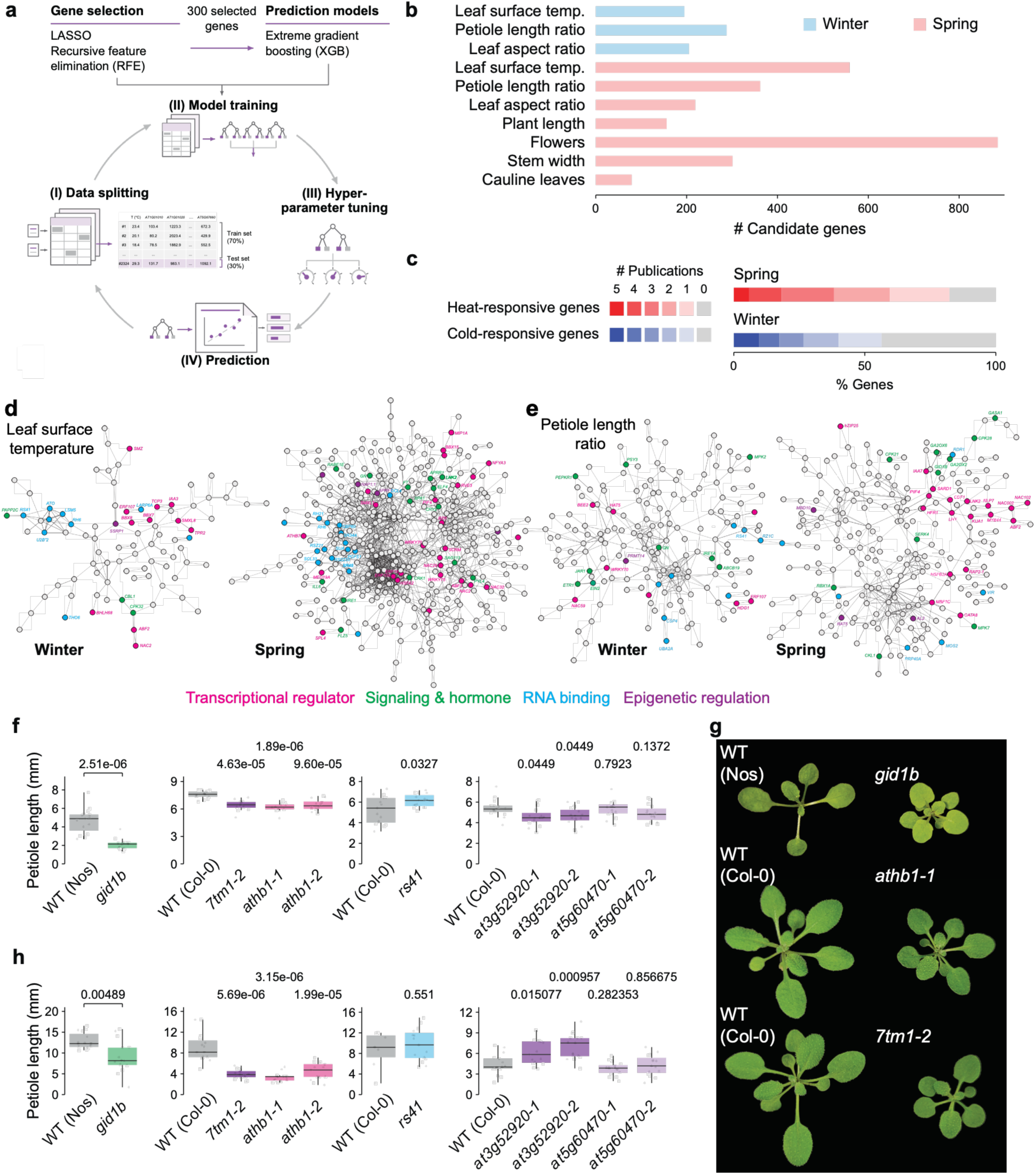
Integrating field transcriptomes and phenotypes with machine learning identifies candidate genes underlying trait variation. **(a)** Schematic overview of the machine-learning workflow. Predictive regression models were trained using Extreme Gradient Boosting (XGBoost) to integrate transcriptomic and phenotypic data from winter and spring collections. Feature selection was performed using LASSO and recursive feature elimination (RFE), yielding a subset of 300 informative genes. Model development included data splitting, training, hyperparameter tuning, and prediction optimization. All models were trained within the season (winter or spring) to exploit inter-individual variation rather than the winter–spring contrast. **(b)** Number of candidate regulators identified by prediction models for each trait. Bar plot showing the number of unique candidate genes identified by models trained on different traits, with independent bars for each trait. Blue and red bars represent winter and spring collections, respectively. In total, 688 and 2,886 candidates were identified for the winter and spring datasets, respectively. **(c)** Validation of model predictions using published temperature-responsive genes. Stacked bars represent the proportion of predicted regulators previously reported to respond to temperature fluctuations in laboratory studies. **(d, e)** STRING gene interaction networks for leaf surface temperature (d) and petiole length ratio (e) candidate regulators in winter and spring collections (STRING v11.5)^43^. Genes with known functions and plausible candidates are highlighted. Disconnected genes are not shown. **(f-h)** Petiole lengths of wild-type plants (Col-0 or Nossen) and selected mutant lines for genes predicted to be involved in petiole length based on the machine-learning approach shown in (a). Plants were grown for 30 days on soil under long-day conditions at 21°C (f, g) and 28°C (h). Statistical significance was assessed using the one-sided *t*-test with unequal variance (*p* < 0.05). Representative WT plants and *gid1b*, *7tm1-2*, and *athb1-1* mutants grown at 21 °C are shown in (g).

We first applied our models to leaf surface temperature, measured purely in situ under natural conditions, to identify candidate genes that predict leaf surface temperature from gene expression. As a direct readout of leaf-level thermal exposure, leaf surface temperature is closely linked to auxin-dependent hyponastic responses that reshape rosette architecture, promote evaporative cooling, and buffer photosynthesis under heat stress^41^. We therefore modeled winter and spring separately (Data S10-11), exploiting within-season temperature variation rather than differences between winter and spring. Using natural variation in leaf surface temperature as an integrative proxy for temperature-dependent transcriptomic states, our models identified 195 candidate genes predictive of leaf surface temperature in winter (1-8 °C) and 559 in spring (6-49 °C), with the highest temperatures likely reflecting transient sun exposure (Fig. 4b; Data S12-13). Three XGBoost models were trained: FULL-XGB, LASSO-XGB, and RFE-XGB (see Methods). All models showed consistent predictive performance across the observed range of leaf surface temperatures in both seasons; however, we further compared the models using root mean squared error (RMSE) and R-squared (R^2^) (Data S15-16). Both LASSO-XGB and RFE-XGB had comparable performance, with median RMSE of 0.947 °C and 0.987 °C and R^2^ of 0.392 and 0.341, respectively, across 10 cross-validation folds in the winter collection (Data S14, Fig S19). In contrast, for the spring collection, RFE-XGB performed best, with the lowest median RMSE and the highest R2 of 4.431 °C and 0.487, respectively (Data S15 and Fig S20). Supporting the relevance of our approach, a substantial fraction of candidates (53% in winter and 85% in spring) have been experimentally shown to respond to temperature shifts under laboratory conditions (Fig. 4c).

Because leaf surface temperature integrates environmental fluctuations in real time, these candidates are expected to include central regulators of temperature-dependent transcriptional programs. Consistent with this interpretation, our predictions recovered well-established regulators of temperature responses, which cluster in a STRING network analysis^42,43^ (Fig. 4d, e). In spring, these included known components of temperature and heat-stress signaling pathways such as the transcription regulators HEAT SHOCK TRANSCRIPTION FACTOR A2 (HSFA2)^44,45^ and EARLY FLOWERING 4 (ELF4)^46^, as well as canonical thermomorphogenesis factors including PIF4^36,47^, which has previously been shown to control leaf surface temperature^48^, CRYPTOCHROME 2 (CRY2)^49^, BRASSINAZOLE RESISTANT 1 (BZR1)^50^, SUPPRESSOR OF PHYA 1 (SPA1)^51^, and the direct PIF4 target FLOWERING LOCUS T (FT)^47^ (Fig. 4d, Data S13). Beyond these established components, our spring predictions highlighted additional transcription factors and hormone-associated regulators, including the ABA receptor PYL4, ABA-RESPONSIVE ELEMENT BINDING FACTOR (ABF3), SUPPRESSOR OF MAX2-LIKE7 (SMXL7), ETHYLENE RESPONSE FACTORs (ERFs), and SCREAM/ICE1, a known regulator of stomatal development. Although many genes have not been directly linked to temperature responses, their regulatory functions make them plausible candidates for integrating thermal cues with hormonal signaling pathways (Fig. 4d, Data S13). In addition, the predicted gene set also contained canonical heat shock proteins, representing established downstream responders to elevated temperatures and further supporting the physiological relevance of our approach (Data S13). In winter, predicted candidates included regulators with established or emerging roles in temperature-related processes, including the calcium sensor CALCINEURIN B-LIKE PROTEIN 1 (CBL1)^52^, TOPLESS-RELATED 2 (TPR2)^53^, PHAVOLUTA (PHV)^54^, and PIN-LIKES 6 (PILS6)^55^ (Fig. 4d, Data S12). Winter candidates also comprised transcription factors, including several members of the B-BOX PROTEIN (BBX) family, SCHLAFMUTZE (SMZ), ABF2, and several RNA binding proteins, which might act as potential regulators (Fig. 4d). Together, these results position our models as a hypothesis-generating framework for identifying candidate genes predictive of temperature-dependent regulatory programs in natural environments. Alongside temperature-dependent responses, circadian regulation and daily environmental cycles also shape short-term variation in the transcriptome. Although sampling was largely restricted to a nine-hour window each day, classification ML predictions were made using collection time of 3 hours windows as traits (Fig S23a). Our ML pipeline showed that using the expression profiles of wild plants to predict collection time yield higher accuracy for samples collected between earlier hours 8:30-11:29 (accuracy = 0.8802) and late hours 14:30-17:30 (accuracy = 0.8629), compared to the samples collected midday (Fig. S23 b, c). Notably our classifier identified multiple regulators involved in circadian processes (Fig S23d, Data S16). These included the clock-associated regulators FLAVIN BINDING, KELCH REPEAT, F-BOX1 (FKF1)^56^, NIGHT LIGHT–INDUCIBLE AND CLOCK-REGULATED 3 (LNK3)^57^, REVEILLE (RVE3)^58^, BBX29^59^ and BBX32^60^, together with regulators of photoperiodic, diurnal and post-transcriptional outputs, including CYCLING DOF FACTOR 2 (CDF2)^61^, TANDEM ZINC KNUCKLE/PLUS3 (TZP)^62^, BBX31^63^ and GLYCINE-RICH PROTEIN 8 (GRP8)^64^ (Data S16).

We next modeled petiole length ratio (PLR), a measure of leaf morphology that integrates growth over time. In total, our models predicted 288 candidate genes associated with PLR in winter and 362 in spring (Data S12 and S13). LASSO-XGB (RMSE = 0.061) and RFE-XGB (RMSE = 0.175) were the best performing PLR prediction models for winter and spring plants, respectively (Data S14 and S15, Fig S21-22). Among the predicted candidates were well-characterized regulators of petiole length, some of which act in a temperature- or light-dependent manner. Notable examples include the transcriptional regulators PIF4 and LONG HYPOCOTYL IN FAR-RED 1 (HFR1)^65,66^, as well as the splicing factors REDUCED RED-LIGHT RESPONSES IN CRY1CRY2 BACKGROUND 1 (RRC1)^67,68^ and ARGININE/SERINE-RICH SPLICING FACTOR 41 (RS41)^68^ (Fig. 4e, Data S12 and 13). Our analysis also identified hormone perception components, including the ethylene receptor ETHYLENE RESPONSE 1 (ETR1) and the gibberellin receptor GA INSENSITIVE DWARF 1B (GID1B) (Fig. 4e). Additional predictors included ETHYLENE INSENSITIVE 2 (EIN2), JASMONATE RESISTANT 1 (JAR1), GIBBERELLIN 2-OXIDASES (GA2OXs), INDOLE-3-ACETIC ACID 7 (IAA7) and IAA26, consistent with contributions from multiple hormonal pathways to petiole growth in the field (Data S12 and S13).

To assess the functional relevance of selected candidates, we examined petiole growth in corresponding mutants under controlled conditions. We selected several of the top-ranked candidates spanning different cellular processes. These included the hormone receptors ETR1^69^ and GID1B^70^; the transcriptional regulators HOMEOBOX1 (AtHB1)^71^ and a DNA-binding protein of unknown function (AT5G60470); SEVEN TRANSMEMBRANE DOMAIN-CONTAINING PROTEIN 1 (7TM1)^72^, which participates in cellulose synthase trafficking; and AT3G52920, which encodes a poorly characterized DUF662-domain protein. Laboratory experiments capture only a subset of the environmental conditions and interactions plants experience in nature, and exhaustively testing these combinations is impractical. As an initial assessment of candidate gene function, we therefore phenotyped the selected mutants grown on soil under standard long-day (LD) conditions at 21 °C. Mutants in *GID1B*, *7TM1*, *AtHB1* and *AT3G52920* exhibited shorter petioles than WT plants, whereas *rs41* mutants showed a modest increase in petiole length (Fig. 4f, g). Mutants in *AT5G60470* showed no detectable difference in petiole length compared with WT (Fig. 4f). When grown at 28 °C, *GID1B*, *7TM1*, and *AtHB1* mutants retained shorter petioles, whereas the *AT3G52920* phenotype was reversed and the *rs41* phenotype was no longer detectable. These observations suggest that growth temperature can modify the phenotypic effects of individual candidates, highlighting the importance of assessing gene function under multiple environmental conditions. To assess temperature-dependent growth responses more broadly, we next performed thermomorphogenesis assays with mutant lines grown on agar plates, quantifying both hypocotyl and petiole length at control and elevated temperatures^73^. These assays revealed genotype × temperature interactions affecting petiole or hypocotyl length in mutants impaired in *ETR1*, *GID1B*, *7TM1* (as well as in the *7tm1 7tm5* double mutant) and *AT5G60470* (Data S17-19, Fig S24-25).

Together, these results support the use of field-derived transcriptomes to prioritize genes involved in growth and temperature responsiveness. The recovery of established regulators, together with functional evidence for additional candidates, illustrates the value of connecting molecular measurements in native populations with experimental validation. Extending this framework across seven measured traits and multiple environments yielded functional predictions for 3,101 genes, providing a resource of testable hypotheses for the Arabidopsis research community (Fig. 4b, Data S14 and S15).

## Discussion

*A. thaliana*, since long being established as a model in plant genetics, is increasingly recognized as a powerful system for dissecting local adaptation to climate change through laboratory studies and common garden experiments. Our work extends this framework by showing that in situ phenotypes and transcriptomes capture climate-associated phenotypic and regulatory variation, providing predictive signatures of gene function under fully natural conditions. This reveals not only the value of *A. thaliana* as a genomic model for climate change studies but also its value as a field-validated system that connects molecular pathways to adaptive potential.

Our observations revisit classical concepts in plant physiology and development, several of which are likely to be crucial for adapting to changing climates. Common garden experiments have shown that *A. thaliana* accessions differ in their compensation after main-shoot damage, with some overcompensating and others undercompensating^35,74^. In our in situ populations, main-shoot loss consistently led to undercompensation in reproductive output, while silique production was shifted towards later developmental stages. At the population level, this temporal spreading of fruit maturation across cooler and warmer periods, observed in Spiekeroog and Brachwitz, is consistent with a bet-hedging strategy^75^. Distributing fertilization and embryo development across heterogeneous thermal regimes may buffer against episodic heat or drought, which are known to negatively affect these processes^76^. Importantly, traits associated with tolerance or resistance to herbivory, although potentially detrimental to individual fitness, may enhance population-level resilience under fluctuating climates by interacting with thermal and hydric stress, underscoring the role of biotic stress responses in climate adaptation. Similarly, early genetic studies uncovered distinct developmental programs controlling longitudinal and lateral leaf growth^77^. Our field observations mirror this distinction: leaf growth prior to winter was predominantly lateral, whereas spring growth was mainly longitudinal, indicating that these developmental programs are plastically deployed across seasons to optimize plant performance under contrasting climatic conditions.

Our results demonstrate that *A. thaliana* phenotypes in situ carry strong, quantifiable imprints of weather, with temperature emerging as the dominant climatic driver of trait variation across years and sites. Redundancy analyses explained 17% of trait variation, with temperature (air, soil, and dew temperatures) accounting for more than 50% of this. While microclimate, edaphic factors, and biotic interactions clearly contribute to the remaining variation, the signal is strong enough to argue that ongoing temperature change will directly reshape *A. thaliana* phenotypes, and, indeed, all plant phenotypes in natural habitats. That a single climatic axis exerts such leverage in fully natural settings is consistent with long-standing field observations that warming is a first-order control on plant form and timing^78,79^, and with the broader insight that experimental warming often underestimates the magnitude of trait shifts observed under real climate anomalies^16^.

Field-derived transcriptomes also carried clear temperature signatures, capturing both immediate responses and integrated environmental histories. Using these signatures, we trained predictive models to identify key regulators shaping transcriptomic and morphological responses to temperature. Recent advances in artificial intelligence and computational modeling have expanded functional prediction^10,30^, but these approaches generally lack ecological and real-world trait data. Our study has started to close this gap. Several genes identified as predictive of phenotypes in the wild are well-known *A. thaliana* regulators previously characterized through laboratory mutant analyses. This highlights that classical regulatory factors also play essential roles under natural conditions, thereby reinforcing the ecological relevance of decades of laboratory research. For example, ETR1, long known for its role in ethylene perception^80–82^, was highly predictive of petiole elongation in the field, and PIF4^36,47^ expression contributed to models of leaf surface temperature and petiole length ratio. Recovering these canonical regulators, along with detecting growth phenotypes in selected candidate mutants, underscores the robustness of our predictive framework. The broader set of candidates represents testable, ecologically grounded functional hypotheses. Natural environments and genetic diversity are only partially captured under standard laboratory conditions; validation across diverse environments and accessions will further refine these predictions. Notably, several previously uncharacterized candidates are associated with epigenetic regulation, mRNA splicing, translation, and metabolic processes, suggesting that environmental responsiveness in the field engages multiple layers of regulatory control. Future landscape transcriptomic approaches incorporating additional molecular layers, such as DNA methylation, alternative splicing, proteomics, and metabolomics, may further enhance predictive resolution and deepen mechanistic insight.

It is projected that the global surface air temperature will increase by 1.5 °C in the near term (2021-2040)^84^. With a growing population and increased per capita exploitation of natural resources, warming events can outpace experimental expectations; functional genomics must increasingly focus on real-world plants under real-world climates. Our study demonstrates that this is both feasible and scientifically transformative: it reconciles laboratory circuitry with field performance and delivers a generalizable blueprint for climate-aware gene discovery. Our field-derived transcriptomes represent regulatory states that integrate both genetic background and environmental inputs, enabling inference of climate-responsive processes without explicitly modeling genotype. Together with large-scale experimental evolution studies such as the GrENE-net project^85^, our results highlight the complementarity between transcriptome-based regulatory inference and population-genomic analyses of climate-driven selection across generations. Nevertheless, our study has limitations, some of which can be addressed in future experiments, whereas others are inherent to observational studies under fully natural conditions. Our observational design does not fully separate environmental, genetic and developmental contributions to expression variation. In particular, our approach cannot fully disentangle environmental, genetic, and developmental contributions to gene expression variation. Future studies incorporating more extensive genomic characterization, fine-scale microclimatic measurements, and additional habitats could help resolve these contributions, although their complete separation remains challenging in natural ecosystems. The complementary survey by Leite Montalvão et al. (2026)^83^ broadened geographic coverage to 60 sites and incorporated microbial profiles, addressing spatial and biotic heterogeneity at the cost of sampling depth within populations. Expression-based predictions and mutant validation support candidate prioritization and functional relevance under the conditions tested, but do not establish direct regulatory relationships or fitness effects in nature. Nevertheless, linking repeated molecular and phenotypic observations captures environmentally associated regulatory states, adding a dynamic dimension to studies of natural variation that cannot be inferred from genome sequence alone. We define landscape transcriptomics as an integrative framework that combines in situ molecular profiling, organismal phenotyping, and quantitative environmental data to infer regulatory architecture directly in natural populations. By treating environmental heterogeneity itself as a systems-level perturbation, this approach enables functional insight without reliance on controlled manipulation. In concert with population-genomic and landscape-scale analyses, landscape transcriptomics offers a path toward identifying key regulators that may enhance the resilience of crops and wild plants to increasingly erratic heat and drought, thereby contributing to ecosystem stability and food security.

## Materials and Methods

### Sample collection sites

3,442 wild *A. thaliana* plants were randomly sampled across two main locations in Germany, Spiekeroog (53.77°N, 7.73°E) and Brachwitz^86^ (51.53°N, 11.87°E) (Figure 1a). Plant samples were collected during their vegetative or reproductive growth stages, which coincide with the local German winter and spring seasons, respectively. Fieldwork was conducted on Spiekeroog between 2021 and 2025, including a winter collection in 2021. At Brachwitz, our samples span 2023-2025 for the spring season only.

### On-site plant phenotyping

For the winter samples, clear aerial images of the sampled plants were taken, and ImageJ^87^ was used to phenotype leaf-related traits, including leaf length, leaf width, the petiole length of the longest rosette leaf, and the total number of leaves. Digital callipers (0-150 mm) were used to obtain quantitative measurements for the spring collection. Phenotyped traits included leaf-related traits, plant length, stem diameter, internodium length, number of cauline leaves, flowers, and side shoots. In addition, an infrared thermometer (−50°C to 550°C) was used to measure the surface temperature of the longest leaf. Information about the plant’s location, including longitude and latitude coordinates, altitude, collection time, and date, was extracted from the images. Supplementary Data S1 describes all measured in situ traits and metadata, and Supplementary Data S2 provides the raw dataset for all plants sampled in the winter and spring collections. Additionally, the standard operating procedure (SOP) used in the field for plant phenotyping is attached (Fig. S27).

### Plant cultivation under laboratory condition

Arabidopsis wild-type (WT; Col-0, Ws-2 or Nossen) and mutant lines (*7tm1*, *7tm5*, *7tm1 7tm5*^72^, *athb1-1, athb1-2*^71^, *etr1-3*^69^, *gid1b*^70^, *at5g60470-1* (SALK_044769), *at5g60470-2* (SALK_061422) were used in this study. Seeds were surface-sterilized, stratified, and plated onto ATS medium supplemented with 1% (w/v) sucrose. Plates were grown vertically in controlled growth cabinets under long-day or short-day conditions with white light at an intensity of 90 µmol m⁻² s⁻¹. Specifically, long day was set as 12h Lights on/12h off for Nossen and *gid1b* while 16h Lights on/8h off for all others and short day was 8h Lights on/16h off for all genotypes. Following germination for 4 d at 20 °C, plates were transferred to either 20 °C or 28 °C and grown for an additional 4 d. Plates were then photographed using a digital camera, and hypocotyl and petiole lengths were quantified from the resulting images. The experimental setup and trait phenotyping followed the single time point analyses described in Janitza et al. (2024)^73^. All the measurements are provided in Data S18-19.

For soil-grown plants, seeds were stratified in 0.1% (w/v) agarose for 4 d and then transferred to soil. Plants were grown under long-day conditions (16 h light, 160-180 µmol m⁻² s⁻¹ / 8 h dark) in a Percival AR-66L3 climate chamber. Total rosette diameter and petiole length of the longest leaf were measured after 30 days. All plants were assigned unique identifiers and phenotyped independently by two investigators in separate measurement rounds, with observers blinded to genotype. Data S17 provides the recorded leaf measurements at 21 °C and 28 °C.

### Plant material, DNA and RNA extraction, RNA-seq library preparation and sequencing

For each spring-measured plant, the longest rosette leaf was collected using forceps and stored in an RNA-preserving solution (25 mM Sodium Citrate, 10 mM EDTA, 700 g/L ammonium sulfate) to preserve RNA integrity for later isolation for up to 5 days. For winter-collected plants, we harvested all aerial parts. After removing excess RNA-preserving solution, plant material was stored at −80°C. RNA and DNA were isolated using the RNA/DNA/Protein Purification High Throughput Plus Kit (Norgen Biotek, Canada) according to the manufacturer’s instructions. A total of 50 ng RNA was used for RNA-seq library preparation using the BRB-seq protocol with some minor modifications^88^. Specifically, a two-step consecutive PCR was introduced to incorporate P5/P7 UDI adapters compatible with Illumina’s NovaSeq platform. For population diversity analyses, 87 samples representing different collection seasons, locations, and years were chosen for Low-Pass Whole Genome Sequencing (LP-WGS) (Data S20). All the RNA libraries were sequenced at Azenta/Genewiz (Leipzig, Germany) on an Illumina NovaSeq sequencer with a 20% PhiX spike-in, and the DNA samples were sequenced at NOVOGENE (Munich, Germany) also on an Illumina NovaSeq sequencer.

### DNA sequence alignment and SNP calling

LP-WGS data was quality-checked, and lower-quality bases (quality score < 20) from the paired-end DNA reads were filtered using FastQC (version 0.12.1; available at https://github.com/FastQC) and TrimGalore (version 0.6.10; available at https://github.com/TrimGalore)^89^, respectively. Reads were aligned to the *A. thaliana* reference (TAIR10) by BWA-MEM with default parameters^90,91^. Samtools sort (version 1.18.0)^92^ was used to sort the resulting alignment file by position, and PCR duplicates in the resulting alignment files were marked by the MarkDuplicates function of GATK (version 4.3.0.0, https://github.com/broadinstitute/gatk).

Single-nucleotide polymorphism (SNP) identification involved estimating the genotype likelihoods, which was done using bcftools mpileup (version 1.18)^93^, restricting analyses to sites with base quality > 20 (BQ > 20) and mapping quality > 20 (MQ > 20). 5903953 raw variants were subsequently called by the bcftools call function. To retain high-confidence SNPs shared across winter and spring collections, we filtered for biallelic sites present in >95% of individuals, with a minor allele frequency ≥ 0.05, variant quality > 50 (QUAL > 50), and mean sequencing depth ≥ 15 (DP ≥ 15); individuals with more than 20% missing SNP genotypes were removed. Filtering was performed using VCFtools (version 0.1.16)^94^. Overall, 78 plant samples and 6054 SNPs were used for downstream diversity analyses.

### RNA sequence alignments, SNP calling, and gene counting

The raw BRB-seq sequencing data consist of two FASTQ files (R1 and R2). R1 contained the P5_BRB_alt2 primer and the different 96 barcode-UMI combinations required for demultiplexing, whereas R2 contained the samples’ RNA fragments. A custom Python script was used to preprocess the R1 file, retaining only sample barcodes (six bases) and UMI (ten bases). The demultiplex function in the BRBseqTools suite (version 1.6.0), with the arguments -p BU -UMI 10, was used to demultiplex reads in the R2 file to their respective plant samples (available at https://github.com/BRB-seqTools)^88^. FastQC (version 0.12.1; available at https://github.com/FastQC) and MultiQC (version 1.19.0) were used for base quality inspection and visualization of the raw demultiplexed reads. Reads from all samples were trimmed to 55 bases using the --hardtrim5 option of Trim Galore; Illumina adapters and low-quality bases (Q < 20) were removed using Cutadapt within Trim Galore (version 0.6.10; available at https://github.com/TrimGalore) ^89^. Filtered reads were aligned to the *A. thaliana* reference genome (TAIR10) using the STAR aligner (version 2.7.11b)^95^ in two-pass mode with arguments -- twopassMode Basic --sjdbScore 2 --outFilterMultimapNmax 15 --outFilterMatchNmin 20 -- outFilterMatchNminOverLread 0.33 --outFilterScoreMinOverLread 0.33. Gene expression quantification was performed using FeatureCounts (version 1.5.3)^96^. Additionally, SNPs were called for selected transcriptomes using the same steps as those previously described for the DNA reads, and PCA analyses were performed for a subset of transcriptomes.

### Population structure and diversity analyses

Population clustering analyses included PCA, phylogeny, and admixture. Prior to the analyses, PLINK (version 2.0.0, https://github.com/chrchang/plink-ng)^97^ was used to process the variants and prune highly correlated SNPs using a sliding window of 50 SNPs and a correlation threshold of 0.1. Additionally, PLINK was used to compute the PCs. A phylogenetic tree was used to infer the evolutionary relationships among plant samples. The variants were re-aligned using MAFFT (version 7.526, https://gitlab.com/sysimm/mafft)^98^ and the gaps were trimmed using TrimAI (version 1.5.1, https://github.com/inab/trimal)^99^. A maximum likelihood (ML) tree was inferred using IQ-TREE (version 3.1.3, https://iqtree.github.io/)^100,101^ and annotated in FigTree (version 1.5.0, https://tree.bio.ed.ac.uk/software/figtree/). To assess genetic differentiation between Brachwitz and Spiekeroog, we computed the pairwise fixation index (Fst). The Weir and Cockerham’s method was used as implemented by the “--weir-fst-pop” function in VCFtools^102^. Furthermore, genetic diversity (□) was also estimated for each population.

### Collection map and in situ trait measurement clustering analyses

Unless otherwise stated, all statistical analyses and visualizations were performed using R (http://www.r-project.org/). Figures were primarily generated using ggplot2 (version 3.5.1). Data preprocessing involved extracting the Brachwitz and Spiekeroog collections across years, standardizing data formats, and merging all the phenotypic data. Geographic coordinates (latitude and longitude) were rendered as a German map using the rnaturalearth (version 1.1.0, available at https://github.com/ropensci/rnaturalearth)^103^ and sf (version 1.0.21, available at https://github.com/r-spatial/sf/)^104^ R packages. Principal component analysis (PCA) was conducted on quantitative traits for winter and spring collections separately using FactoMineR (version 2.12.0, available at https://github.com/husson/FactoMineR)^105^. To evaluate the effects of season, year, and location on plant clustering based on leaf-related traits, we applied Factor Analysis of Mixed Data (FAMD)^105^, which integrates quantitative and categorical variables.

### Statistical analyses

Assumptions of normality and homogeneity of variance between groups (seasons or years) were assessed using Shapiro-Wilk and Levene’s tests, respectively. As most traits deviated from normality (Shapiro-Wilk test, *p* < 0.05; Supplementary Data 16), non-parametric methods for group comparisons were used. Pairwise comparisons were performed using the Wilcoxon rank-sum test across years, locations, and between main shoot yes/no groups, while the Kruskal-Wallis test was applied for comparisons involving more than two groups. Unless stated otherwise, all tests were two-sided, and statistical significance was defined as p-value (*p*) < 0.05.

### Seasonal influence on leaf-related in situ trait measurements

Multivariate analysis of variance (MANOVA) was performed to assess the influence of seasonal shift on leaf-related traits. For this, the plants collected at Spiekeroog in 2021 (winter, n = 684) were compared with those collected in the same areas in the corresponding spring (n = 125) of 2022. To identify significant differences between traits, we performed one-way univariate ANOVA tests on the MANOVA results as a post hoc analysis. The results were visualized as boxplots with p-values. Additionally, individual Spearman correlations were calculated for each collection to understand the dynamics in the leaf-related traits relationship.

### Yearly and location comparisons of in situ measurements collected during spring season

Spring collections span different years across both locations and covered more traits, thus allowing more comparisons. To assess overall trait associations while accounting for hierarchical sampling and environmental structure, linear mixed-effects models were fitted to evaluate relationships between all possible pairs of spring trait measurements. Models were implemented and fitted using lme4 (version 1.1-37, available at https://github.com/lme4/lme4/)^106^ with year and location specified as random intercepts and traits as predictors and response variables. All traits were standardized prior to modeling to facilitate comparability of model effect estimates. Model coefficients and p-values were extracted from each pairwise model, and p-values were adjusted for multiple testing using the Benjamini-Hochberg (BH)^107^ procedure. Effect estimates were visualized as a correlation heatmap using the ggcorrplot R package (version 0.1.4.1, available at https://github.com/kassambara/ggcorrplot). Pairwise trait correlations were assessed using Spearman’s rank correlation for individual season-year combinations for each location.

To further investigate trait variation across years at each location, we performed a non-parametric (Kruskal-Wallis) test. Lastly, reproductive growth phase traits that are affected by the loss of the main shoot, mainly the number of flowers, siliques, side branches, and side shoots, were analyzed. Additionally, reproductive output, which is the sum of flowers and siliques on a plant, was generated. The comparison between plants with a main shoot (Y) and without (N) for these traits was performed using the Wilcoxon rank-sum test.

### The effect of weather variables on trait variation

Daily weather data at 10km resolution for Spiekeroog (2020-2025) and Brachwitz (2022-2025) were retrieved from the open-source Open-Meteo (https://open-meteo.com/; Supplementary Data 3). Monthly means were calculated for wind speed, relative humidity, dew point, sunshine duration, air and soil temperatures, while daily precipitation values were aggregated to monthly totals. To enable direct comparison of interannual fluctuations in weather variables, relative anomalies were calculated for each weather variable as the difference between the observed monthly mean and the corresponding monthly baseline mean for 2021-2025 and 2022-2025 at Spiekeroog and Brachwitz, respectively. Monthly baseline means were obtained by averaging each variable across collection years for a given month. To explore whether fluctuations in weather variables were reflected in our traits, the association between the mean air temperature of the four months prior to April and petiole length was tested across the spring collection. Furthermore, the petiole length ratio was used to compare variation in petiole length across years in Brachwitz and Spiekeroog, as this trait could indicate temperature effects^36^.

To extend these analyses, we investigated how environmental conditions relate to all spring trait measurements using linear mixed-effects models. Prior to the analysis, highly correlated weather variables were excluded, and final models included air temperature, precipitation, wind speed, and sunshine duration. The average across December, January, February, and March for the weather variables were fitted as fixed effects, along with traits and the weather-trait interactions. These models allowed us to estimate the magnitude and direction of the weather-trait relationship.

To systematically assess the influence of weather variables on variation across all recorded traits and collection years, we performed redundancy analysis (RDA) using the vegan R package (version 2.8.0) (https://github.com/vegandevs/vegan)^108^. Prior to RDA, significant weather variables (p < 0.01) were identified by forward selection using the adespatial R package (version 0.3.28).

### Data preparation, gene counts normalization, and batch correction

Samples were sequenced across multiple batches. Supplementary Data S21 provides a complete list of plant identifiers and their corresponding library preparation and sequencing batches.

1. For differential gene expression (DGE) analyses for seasons (Spiekeroog, winter vs spring), leaf temperature (lower and upper leaf surface temperature), and location (Spring samples, Spiekeroog vs Brachwitz);

To account for batch effects, count datasets were first separated by season, and experimental biases were removed using the empirical Bayes framework implemented in the Surrogate Variable Analysis (SVA version 3.54.0)^109^ with the Combat-seq function. This approach preserves the integer nature of count values and therefore makes the counts compatible with downstream normalization, specifically for DGE analysis^110^. Lowly expressed genes were removed using a threshold of an average expression below 10 raw counts across all samples. The remaining genes were normalized to counts per million (CPM) using DGEobj.utils (version 1.0.6 available at https://github.com/cran/DGEobj.utils) and edgeR (version 4.6.3, available at https://bioconductor.org/packages/edgeR)^111^. All intermediate datasets, including batch-corrected, filtered, and normalized expression matrices, are available at the link associated with the analyses for this manuscript.

1. For gene selection and gene-trait predictive modeling;

Individuals with more than 95% of zero gene counts were excluded, followed by filtering of lowly expressed genes (sum of gene counts < 10). Count values were normalized within each batch using variance-stabilizing normalization (VSN) implemented via the vst function of DESeq2 (version 1.46.0) with default parameters^112^. Samples were separated by season, and batch correction was performed using a data-adaptive approach (ARSyN) implemented in the NOISeq package (version 2.50.0)^113,114^. This batch-correction strategy effectively removed non-biological sources of variation, such that season of sample collection emerged as the primary axis of transcriptomic divergence (PC1; Fig. S27). Finally, to retain the most informative features for downstream analyses, low-variance genes (below the 80th percentile of variance across all genes) were excluded, and expression values were log₂-transformed.

### Characterization of microbial community from unmapped transcriptome reads

BRB-seq reads that did not align to the *A. thaliana* reference genome were used as input for taxonomic classification using Kaiju (version 1.10.1)^115^. Analyses were performed against the nr_euk database, which comprises protein sequences from archaea, bacteria, fungi, microbial eukaryotes, and viruses. Default parameters were used, except that the maximum number of allowed mismatches was set to five and the minimum match length was set to 50. Resulting taxonomic profiles were formatted and analyzed using the R packages phyloseq (version 1.48.0)^116^ and vegan (version 2.7.1)^108^.

### Differential gene expression and GO analysis

Considering the unusually large sample sizes across seasons and locations in this study, conventional parametric approaches for differential expression analysis, such as those implemented in DESeq2^112^ and edgeR^111^, were not optimal. Instead, a non-parametric Wilcoxon rank-sum test was applied to identify differentially expressed genes (DEGs) from matrices of CPM-normalized expression counts between groups, following the approaches described by Li et al. (2022)^117^ and Redmond et al. (2024)^118^. Differential expression analyses comparing winter versus spring samples (Spiekeroog only) and Spiekeroog versus Brachwitz samples (restricted to 2023 and 2024, when both locations were sampled) were referred to as seasonal and locational DEG analyses, respectively.

To identify DEGs associated with leaf surface temperature, plants were stratified based on the distribution of their measured temperatures. Specifically, the lower and upper 10^th^ percentiles were calculated, and plants below the 10^th^ percentile and above the 90^th^ percentile were classified as low- and high-temperature groups, respectively. Resulting p-values were adjusted for multiple testing using the Benjamini-Hochberg (BH)^107^ procedure. Genes were considered significantly differentially expressed if they met both criteria: (i) adjusted p < 0.01 and (ii) |log₂ fold change| > 1.

Gene Ontology (GO) term overrepresentation analysis for biological processes was conducted on the DEG sets from each analysis using the gprofiler2 R package (version 0.2.3)^119^. Multiple-testing correction was performed using the BH method, with a user-defined significance threshold of p < 1 × 10⁻⁴.

### Gene selection

Log_2_-transformed gene counts from all samples (winter and spring separately) were used as input for gene selection using least absolute shrinkage and selection operator (LASSO)^120^ and recursive feature elimination (RFE)^121^. These complementary machine learning approaches were employed to identify the top 300 genes most informative for predicting a trait of interest in our collection (e.g., leaf surface temperature). LASSO regression was implemented using elasticnet (version 1.3)^122^, whereas RFE was performed using the rfe function with a random forest as an external learning algorithm (rfFuncs), as implemented in the caret R package (version 7.0-1, https://github.com/caret/)^123^.

Model training and hyperparameter tuning were carried out using 10-fold cross-validation for each algorithm. The model with the lowest root mean squared error (RMSE) was selected as the best-performing model. Employing both LASSO (linear, sparsity-inducing) and RFE (tree-based, performance-driven) enabled complementary gene selection strategies. Supplementary Data S22 and S23 provides all the selected feature from LASSO and RFE models trained for different traits in winter and spring collections, respectively.

### Predictive modelling of individual traits

Two strategies were used for predictive modeling: (i) regression models trained on subsets of genes selected by LASSO and RFE for each trait, and (ii) models trained using all available gene features. The overall workflow, including data partitioning, model training, optimization, and evaluation, is illustrated in Fig. 4a. Data were split using a 70:30 ratio, with 70% of plants used for training and 30% reserved for testing. Predictive models were constructed using extreme gradient boosting (XGBoost) regression, implemented via the xgbTree function in the caret framework. Hyperparameters, including learning rate (η), maximum tree depth, number of boosting rounds, and subsampling ratio, were optimized using grid search within caret, with 10-fold cross-validation. RMSE was used as the primary metric for model selection.

Winter and spring samples were modeled separately. Models predicting leaf surface temperature, petiole length ratio, and leaf aspect ratio were trained for both seasons, whereas models for plant length, stem width, number of cauline leaves, and flower number were trained using spring samples only. Additionally, we investigated for the genes that their expression are influenced by the circadian rhythms and thus the time of the sample collection. For this, winter plants were used and samples were grouped into bins of three hours (8:30 am - 11:29 am, 11:30 am - 14:29 pm, 14:30 pm - 17:29 pm). A classification model was trained on these three classes using the LASSO algorithm and genes corresponding to each bins were extracted and reviewed.

We highlight the performance of these models for their respective traits, mainly leaf surface temperature and petiole length ratio, across winter and spring samples (Supplementary Figs 20-23). All performance metrics and plots for other traits (leaf aspect ratio, number of cauline leaves, flowers, stem width, and plant length) and the resulting model objects (from R) are provided in the Zenodo script repository associated with the manuscript.

### Comparative validation of temperature-responsive candidate genes

To validate candidate genes associated with leaf surface temperature predictions, a comparative analysis was performed against previously published transcriptomic studies of *A. thaliana* aerial tissues exposed to cold^124–128^ and heat^129–133^ treatments. Raw RNA-seq data were retrieved from NCBI and reprocessed uniformly. All datasets were analyzed using the nf-core RNA-seq pipeline (version 3.18.0)^134^, with read alignment performed using STAR (version 2.7.11b)^95^ and transcript quantification using Salmon (version 1.10.3)^135^.

Differential expression analysis was conducted using DESeq2 (version 1.49.3)^112^, and genes with p < 0.05 and |log₂ fold change| > 1 were considered significantly differentially expressed. Significant DEGs from these studies were compared with candidate genes identified in our predictive models, and overlap significance was assessed using a hypergeometric test. Finally, genes were grouped according to the number of studies in which they overlapped (0-5).

### Protein-protein interaction of candidate genes

The nonredundant gene candidates were extracted from the three XGBoost models for leaf surface temperature and the petiole length ratio, respectively. These were used as input for protein-protein interaction network analyses, which were generated by STRING (version 12.5)^43,136^.

## Data availability

All raw FASTQ files, intermediate next-generation sequencing files, and associated metadata are publicly available in the European Nucleotide Archive (ENA) at EMBL-EBI under project accession PRJEB89549 (https://www.ebi.ac.uk/ena/browser/view/PRJEB89549). Source data necessary for plotting and ML analyses are provided in this paper as supplemental datasets. We retrieved the *A. thaliana* reference genome (TAIR10)^137^ and transcriptome (AtRTD3)^138^ from https://www.arabidopsis.org/ and https://ics.hutton.ac.uk/atRTD/RTD3/, respectively. We performed a comparative analysis of publicly available RNA-seq datasets from the publications mentioned above. Meteorological measurements were freely retrieved from https://open-meteo.com/. Additionally, the supplemental dataset associated with this work can be found at https://doi.org/10.5281/zenodo.23728284.

## Code availability

Scripts for all analyses and the associated datasets are available on Zenodo^139^ at [https://doi.org/10.5281/zenodo.23816872].

## Author contributions

Conceptualization: EYM, MLB, HB, MQ and SL Data analysis: EYM, MLB, CFH, ABW, MKZ Phenotyping: DCA, CA, MLB, HB, VB, BB, MC, LF, SH, ELG, RH, PH-N, TJ, UJ, MJ, ALK, KK, MKl, MKo, CK, SL, EL, SM, FM, EYM, JM, PP, MQ, HR, TR, BR, CS, TS, CRS, LS, FS, RS, AS, DS, DW Transcriptome libraries: MJ, BV Writing: EYM, MLB and SL with contributions from all authors

## Acknowledgments

We are grateful to Heather E. McFarlane, Raquel Chan, Svenja Saile, Andreas Wachter and NASC for providing seeds, and to Christa Lanz for assistance with sequencing. We gratefully acknowledge Wiebke Dammann, Marvin Dill, Nadja Janzen, Johannes Köther, Kevin Krawetzke, Svea Küper, Felix Meyer, Anica Schmauch, Friedrich Schönauer, Sandra Schüler, Phil Weikert, and Joachim Weber for their initial support and contributions to this project. We thank the staff from the Lower Saxony Wadden Sea National Park and the Nationalpark-Haus Wittbülten on Spiekeroog for granting us permission and access to the field, as well as their logistical support and advice. During the preparation of this work, the authors used ChatGPT for language editing, including spelling correction and readability improvements, and q.e.d. for an initial assessment of the manuscript. After using these tools, the authors reviewed and edited the content as needed and take full responsibility for the content of the publication.

## Funding

This research was supported by the Deutsche Forschungsgemeinschaft (DFG-514901783) through the Collaborative Research Centre 1664 “Plant Proteoform Diversity-SNP2Prot” (projects A02 and A05), LA2633/4-2, LA2633/6-1 (all DFG) and the EFRE-funded consortium ValuePlant. Sampling in the dry grasslands near Brachwitz was supported by the FlexPool project iSPOT (Spatial scaling of porphyry outcrop trends), funded by the German Centre for Integrative Biodiversity Research (iDiv) Halle-Jena-Leipzig.

## Competing interests

The authors declare no conflict of interest.

## Notes

### Competing Interest Statement

The authors have declared no competing interest.

### Summary of Updates

This revision adds genomic analyses and an analysis of collection-time-associated gene expression, expands the reporting of predictive models and mutant phenotypes, and clarifies the interpretation of the field observations. We have also expanded the discussion of previous field transcriptomic studies and clarified how our approach combines individual transcriptomes with quantitative phenotyping in naturally established populations. Methods, figures, legends, supplementary information, and data and code availability statements have been updated.

