## Supplemental Figures for "Molecular and phenotypic footprints of climate in native *Arabidopsis thaliana*"

### Supplementary Figures for Mjema et al., 2026

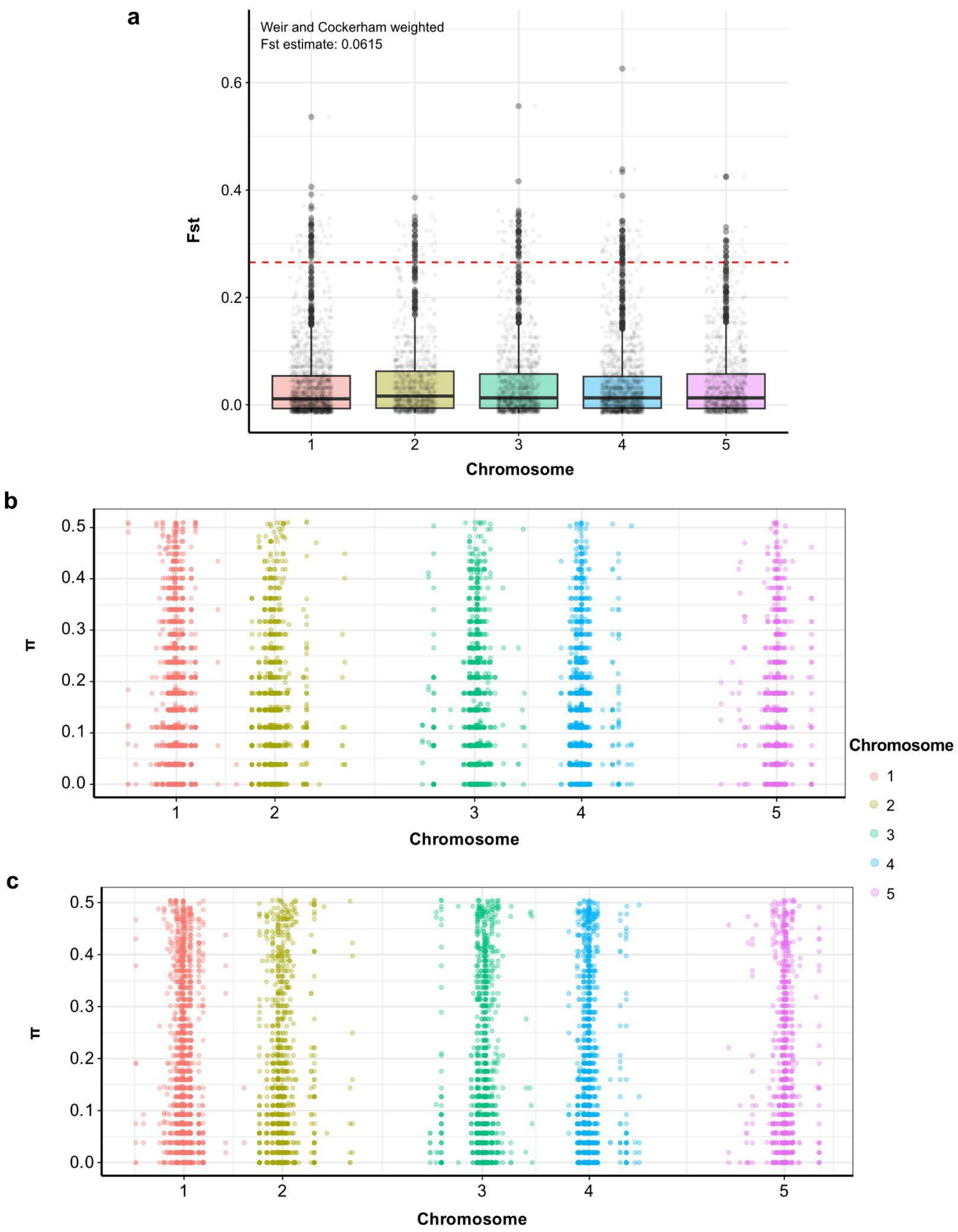


**Fig S1: Genetic differentiation and diversity across the five *A. thaliana* chromosomes.**

**a:** Distribution of fixation index (Fst) on each chromosome. Each point represents an Fst value for individual SNPs (6054). The 97.5th percentile Fst threshold (0.265) is indicated as a red-dotted line above which the SNPs are considered significantly genetically different between Brachwitz and Spiekeroog.
**b-c:** Nucleotide diversity (π) per each SNP position for (b) Brachwitz and (c) Spiekeroog. The mean π for Brachwitz and Spiekeroog were 0.15983 and 0.17442, respectively.

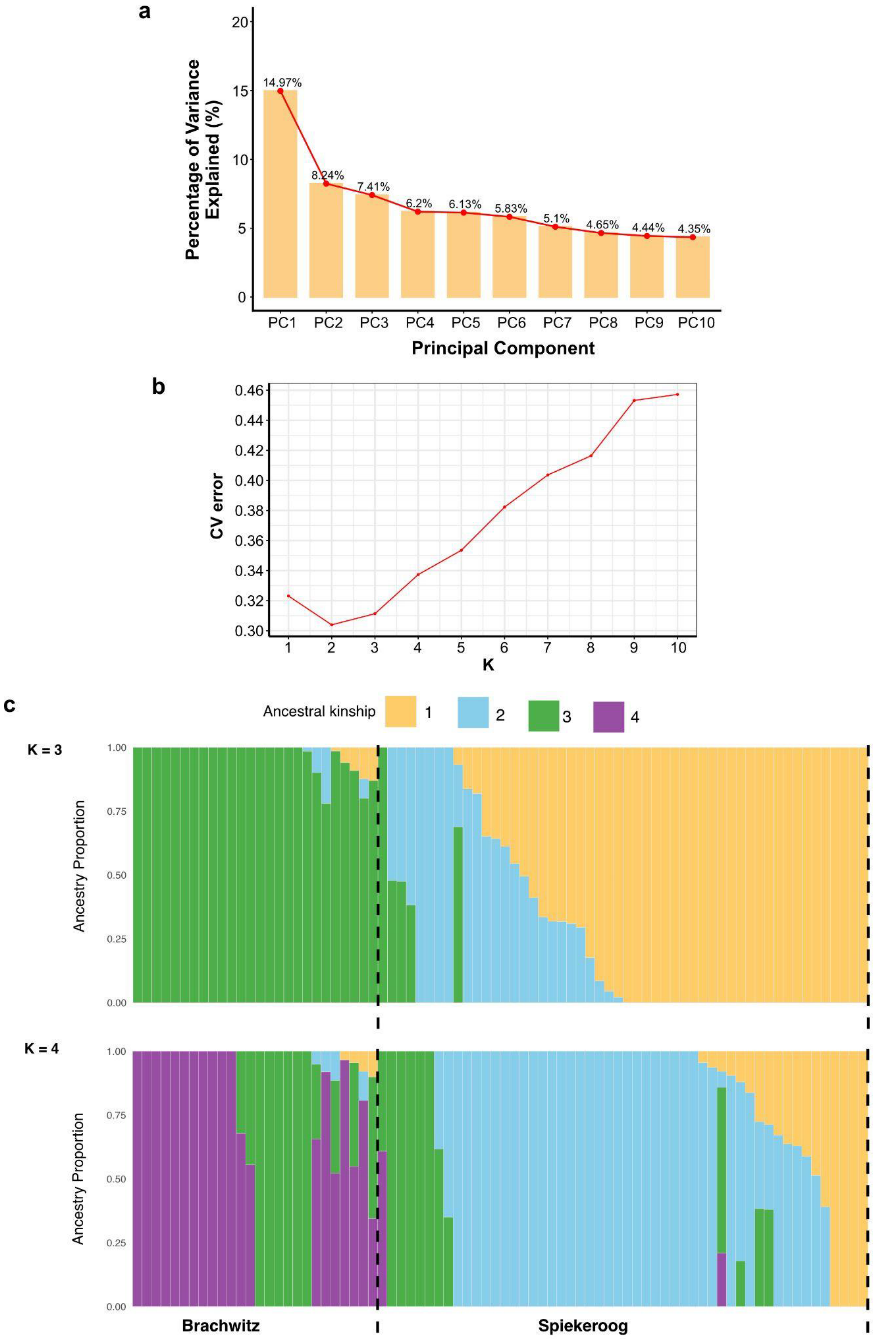


**Fig S2: Genetic relationship between wild *A. thaliana* accessions.**

**a:** A PCA scree plot showing the first 10 PCs and their respective percentage of variation explained.
**b:** Line plot showing the CV error (y-axis) for 10 ancestral kinship tested in the admixture model.
**c:** Admixture bar plots for K = 3 (top panel) and K = 4 (bottom panel). High population admixture and subpopulation appeared at higher K. Each bar represents a plant sample. The bar color and color length indicates the K and its proportion, respective. Each K is coloured differently as indicated on the legend at the top.

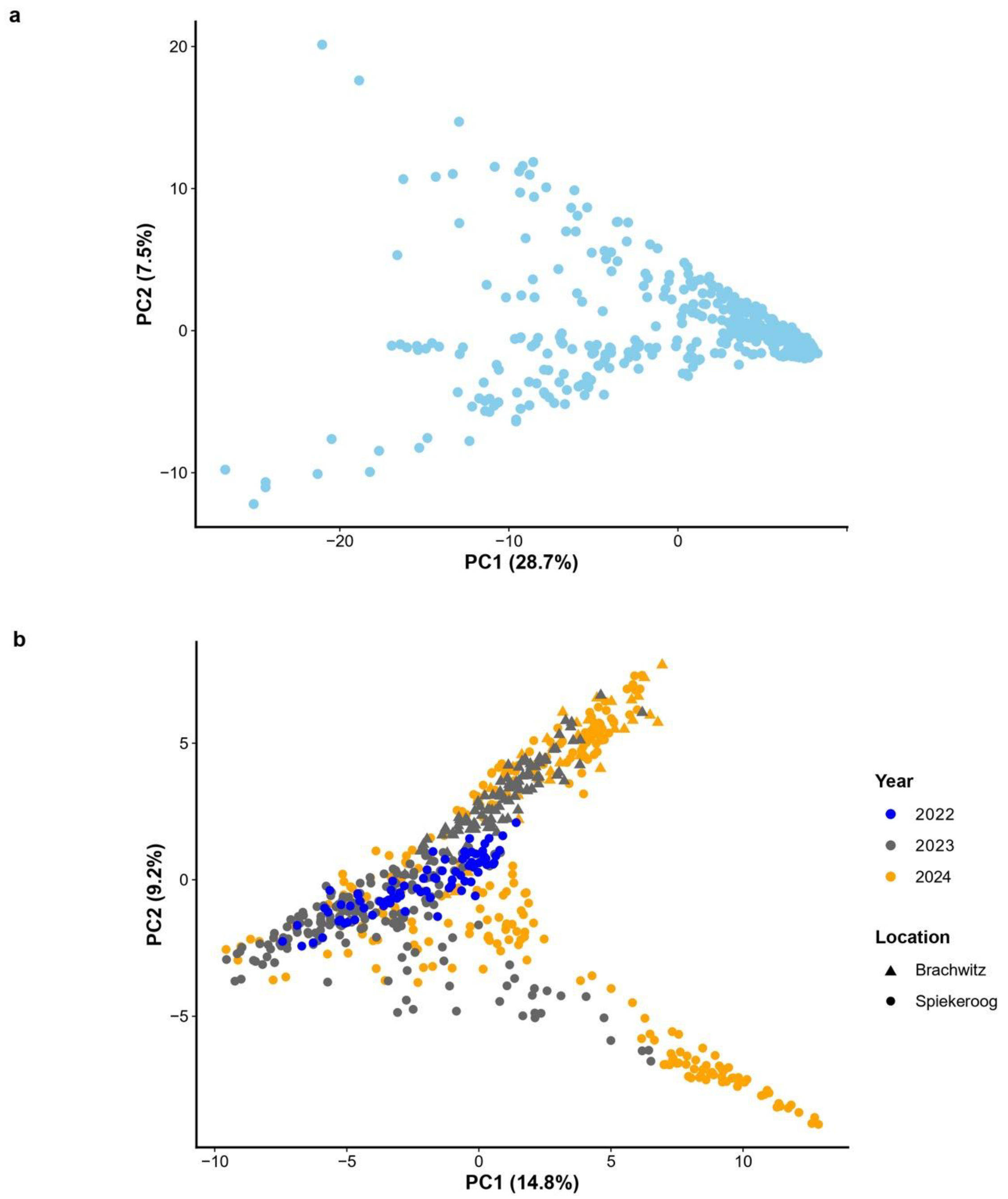


**Fig S3: PCA biplot on SNPs for winter and spring transcriptomes showing continuous pattern of variation with limited genetic clustering among selected plants.**

**a, b:** PCA of transcription-based SNPs from a subset of our collection from (a) winter and (b) spring collections show substantial overlap across years and locations, with PC1 and PC2 explaining a modest proportion of total genetic variance.

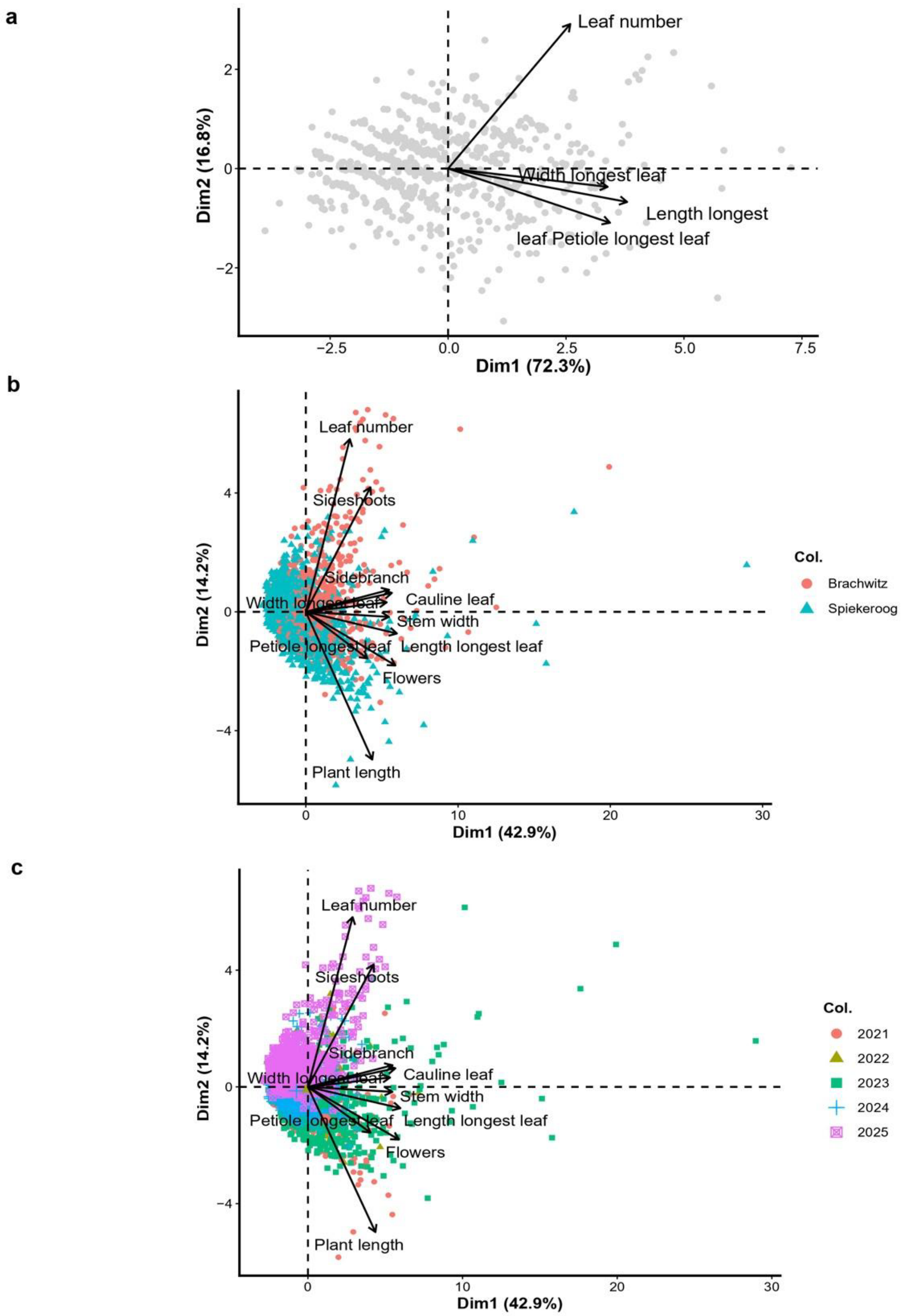


**Fig S4: The PCA biplot based on quantitative trait measurements showed no clear clustering of samples.**

**a-c:** PCA biplot of (a) winter samples, and (b, c) spring samples colored by collection (b) location, and (c) year. No clustering pattern indicative of season or location was observed but rather minimal yearly clustering; hence, the majority of the variability in the clustering analysis stemmed from differences in trait expression.

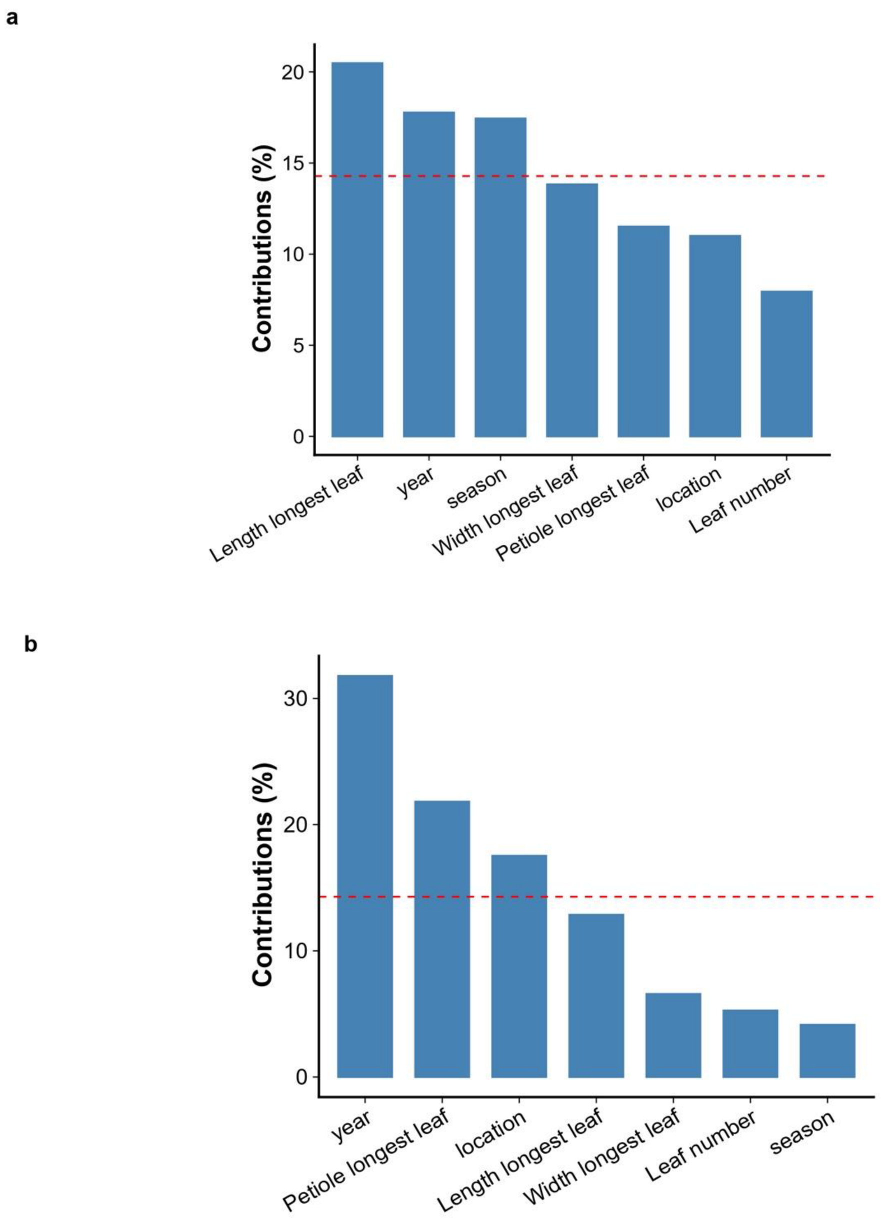


**Fig S5: Leaf length, collection year, and season had the highest contribution in dimensions 1 and 2.**

**a and b:** Bar plot showing the percentage contribution of different variables used in FAMD analysis. Along dimension 1 (a), plant leaf length, year, and location of collection contributed more, as depicted in Figure 1e. The influence of both location and the length of the plant's petiole was depicted on dimension 2 (b), in which the collection year had the highest contribution.

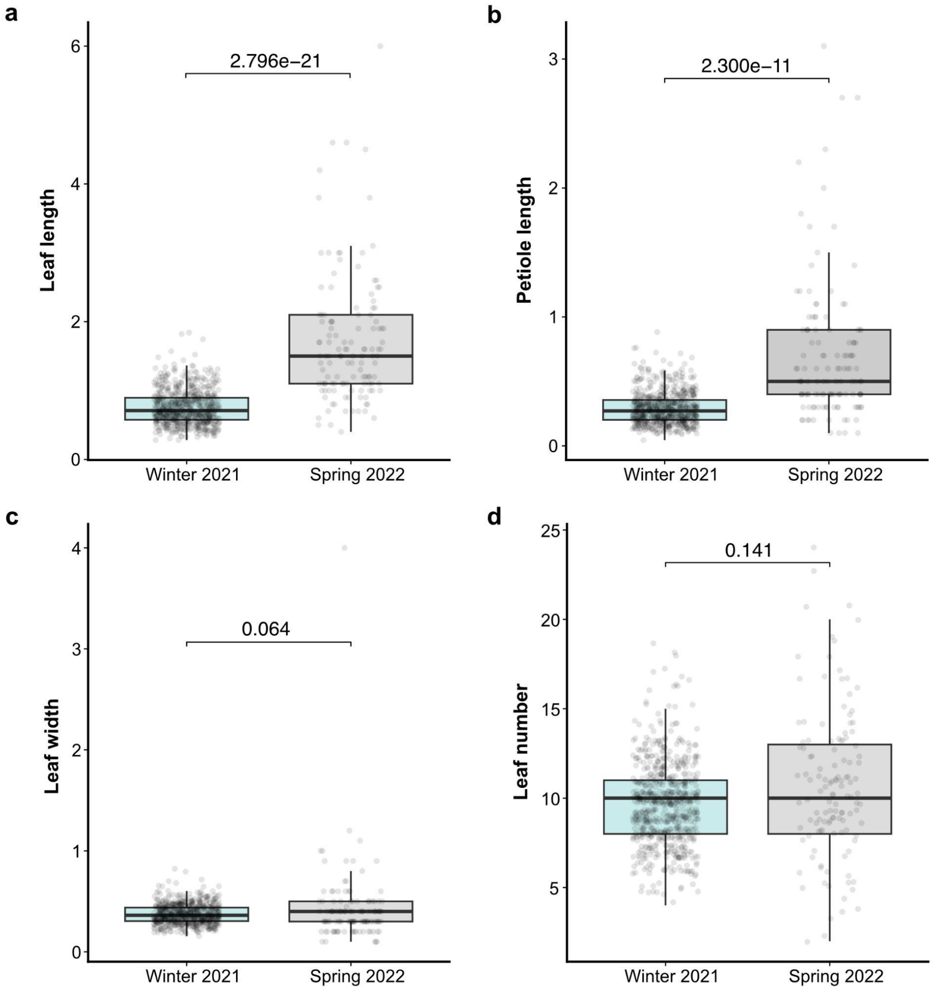


**Fig S6: Seasonal differences in leaf-related traits at Spiekeroog.**

**a-d:** Box plots depicting the comparison between different leaf-related traits between winter (2021) and spring (2022). Leaf and petiole length significantly increased from winter to spring (a, b). No significant differences were observed in leaf width (c) and number of leaves produced (d), suggesting that the main direction of growth is longitudinal rather than lateral during seasonal transitions. One-way univariate ANOVA on the MANOVA result with statistical significance defined as $p < 0.05$.

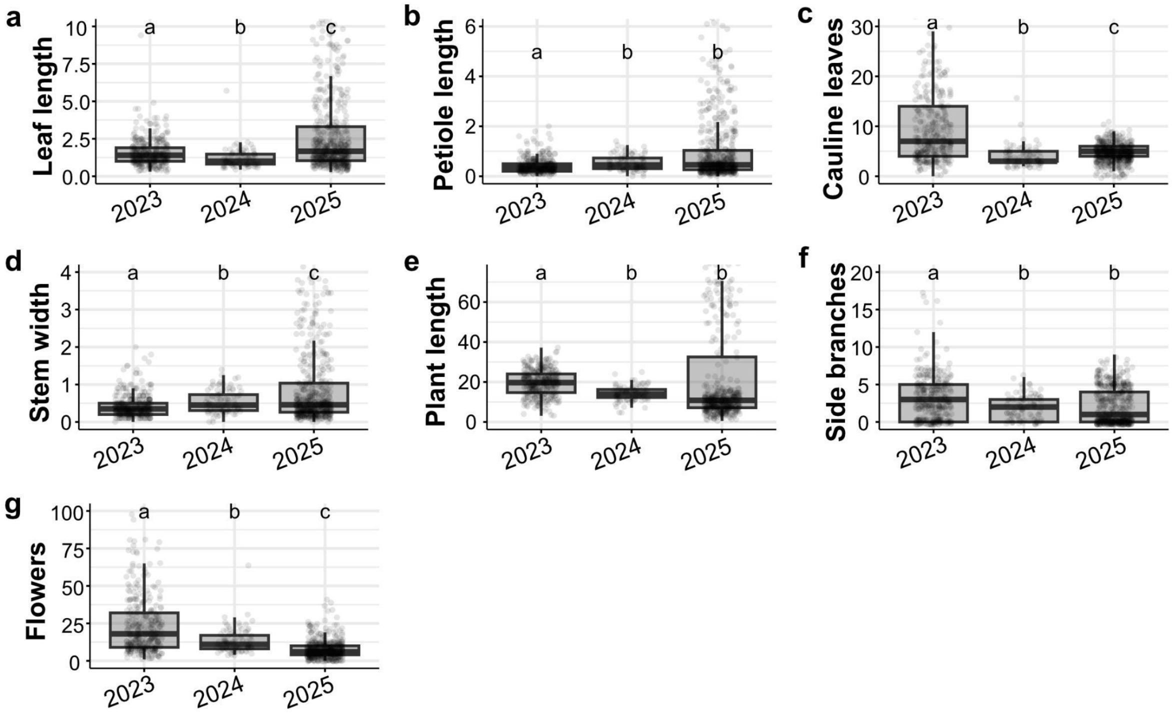


**Fig S7: Phenotypic variation in Brachwitz spring populations (2023-2025).**

**a-g:** Box plots showing various selected trait comparisons for 352, 95, and 564 plants collected in the spring of 2023, 2024, and 2025, respectively. A significant shift in trait variation was observed in at least one comparison pair for both vegetative traits (a-b) and reproductive traits (c-g). Different lowercase letters indicate significant differences ($p < 0.05$) from the Dunn's post hoc test of the Kruskal-Wallis results.

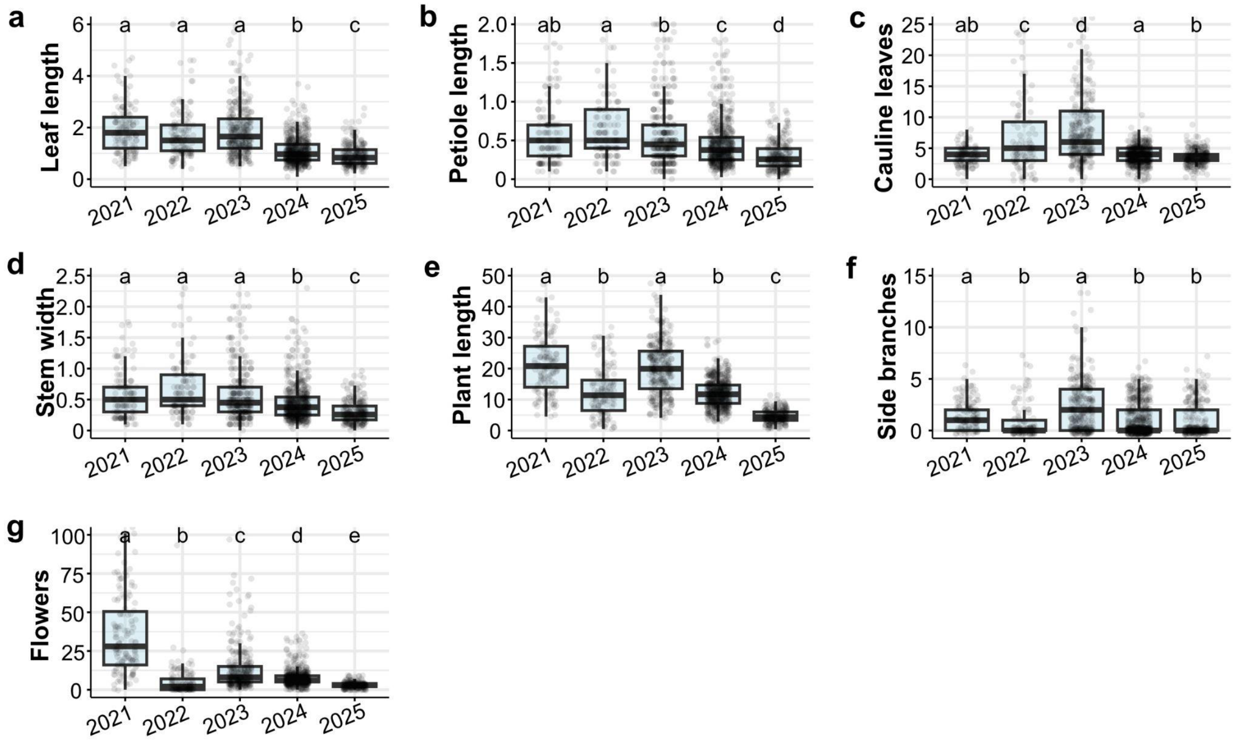


**Fig S8: Phenotypic variation in Spiekeroog spring populations (2021-2025).**

**a-g:** Box plots showing trait comparisons for 130, 125, 302, 557, and 246 plants collected in spring of 2021, 2022, 2023, 2024, and 2025, respectively. A significant shift in trait variation was observed in at least one comparison pair for both vegetative traits (a-b) and reproductive traits (c-g). Different lowercase letters indicate significant differences ($p < 0.05$) from the Dunn's post hoc test of the Kruskal-Wallis results.

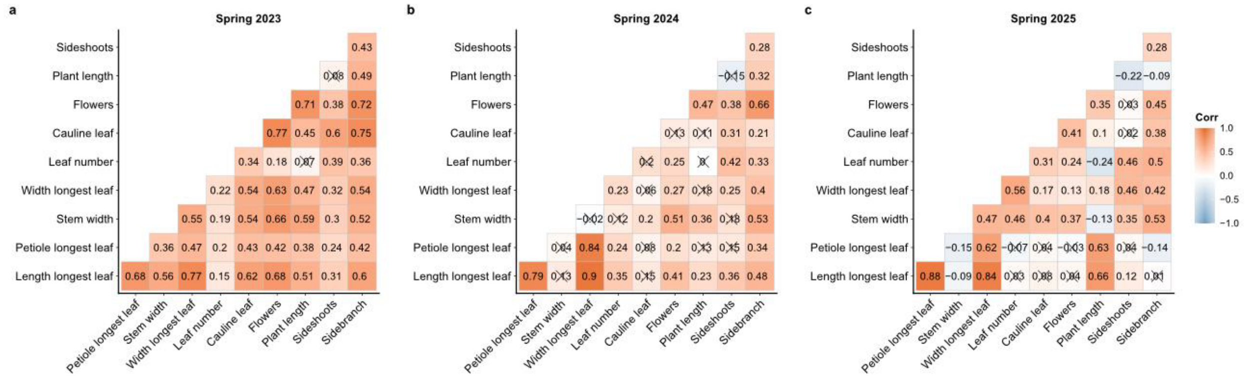


**Fig S9: Correlation heatmap for plants collected from Brachwitz.**

**a-c:** Heatmaps displaying Spearman correlation between trait measurements for (a) 2023, (b) 2024, and (c) 2025 spring plants in Brachwitz, respectively. Tile colours indicate the strength and direction of the trait relationships. The "x" sign indicates non-significant relationships (statistical significance was defined as $p < 0.05$).

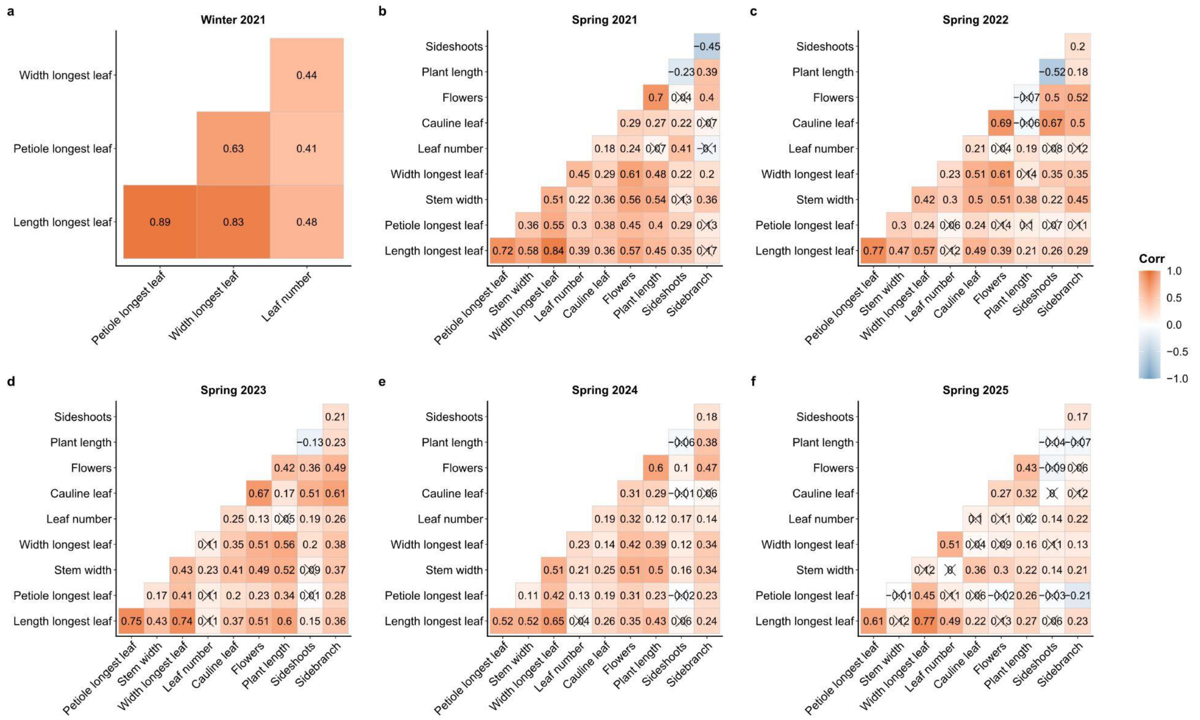


**Fig S10: Correlation heatmap for plants collected from Spiekeroog.**

**a-f:** Heatmaps displaying Spearman correlation between trait measurements for (a) winter 2021 and (b-f) spring of 2021-2025 plants in Spiekeroog, respectively. Tile colours indicate the strength and direction of the trait relationships. The "x" sign indicates non-significant relationships (statistical significance was defined as $p < 0.05$).

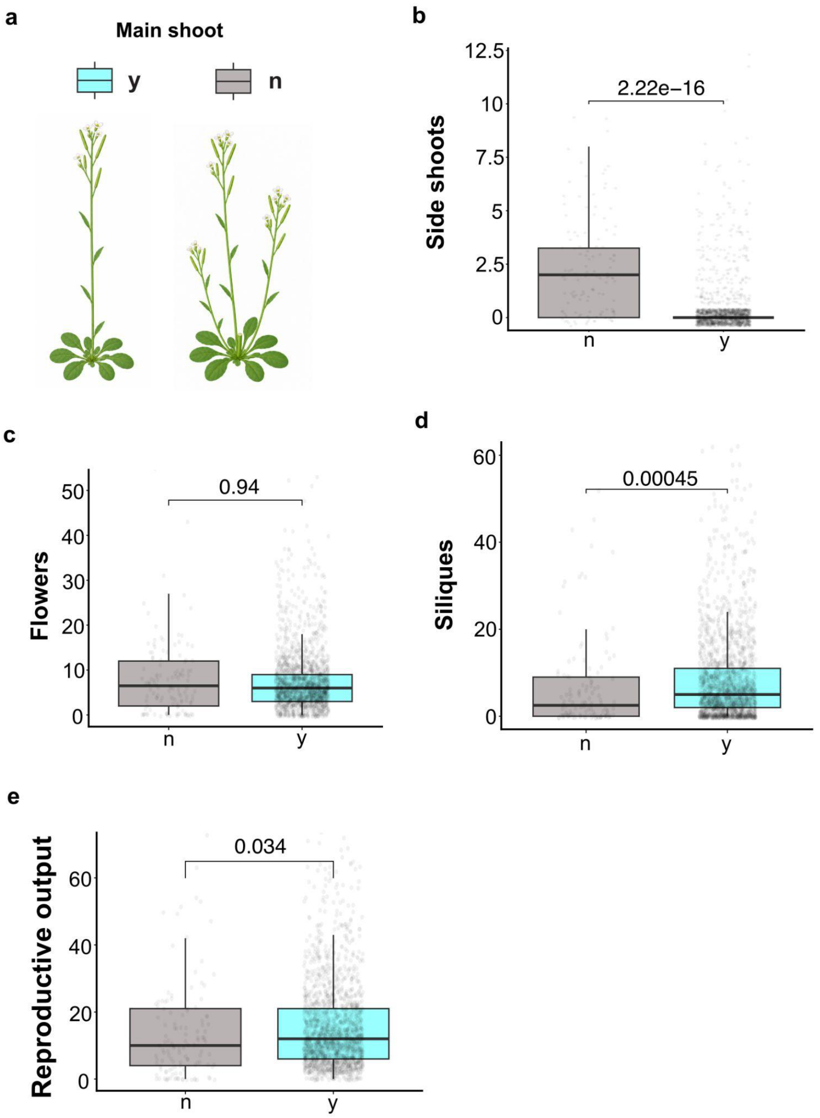


**Fig S11: The presence or absence of the main shoot impacts reproductive output.**

**a:** Schematic representations (top left) illustrate plants with (y) or without (n) a main shoot.
**b-e:** Boxplot showing the comparison between plants with and without a main shoot in the number of (b) side shoots (c) number of flowers (d) number of siliques and (e) reproductive output. There was no significant difference in the number of flowers; however, plants with the main shoot produced more siliques, thereby positively influencing reproductive output. (Wilcoxon rank-sum test with statistical significance defined as $p < 0.05$).

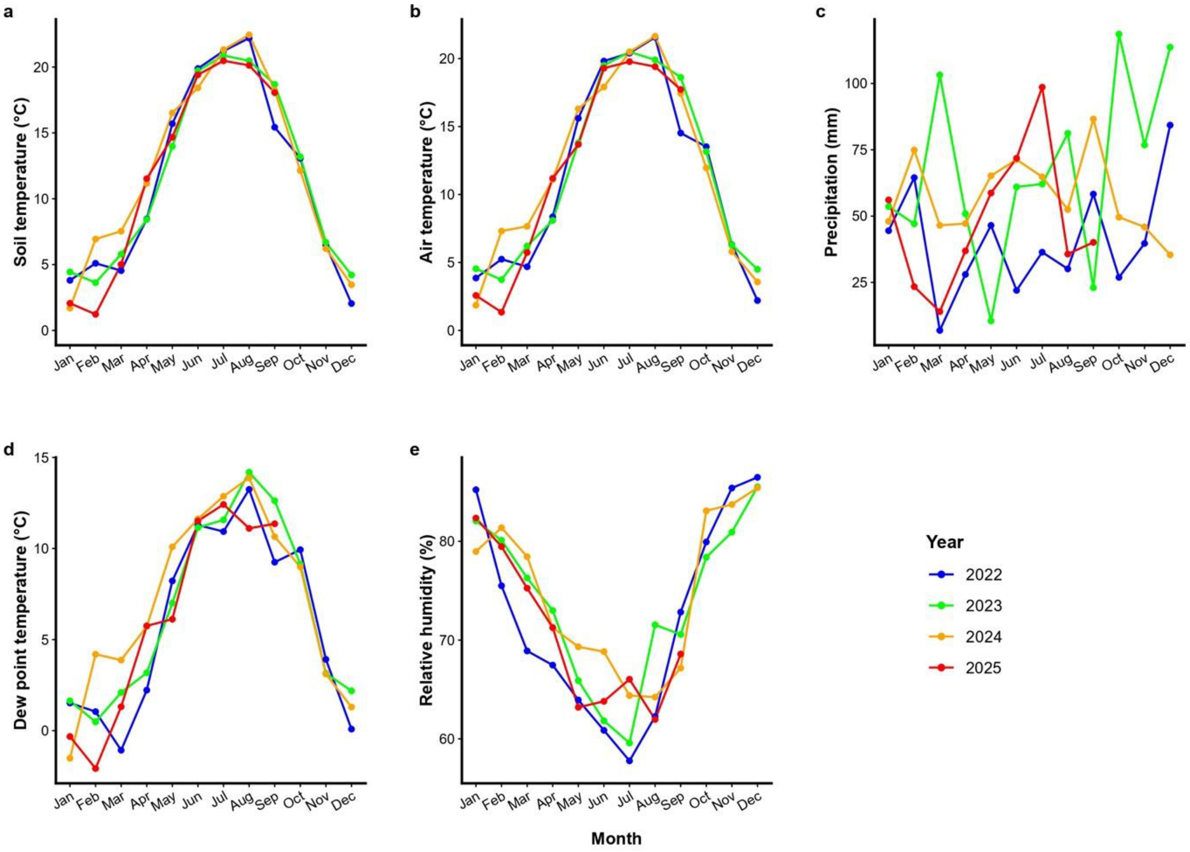


**Fig S12: Interannual variations in the mean monthly weather variables in Brachwitz.**

Line plots of different climatic factors showing trends in monthly changes for each parameter in Brachwitz (2022-2025). Displayed are (a) Soil, (b) Air, and (d) dew point temperatures, (c) precipitation, and (e) relative humidity trends.

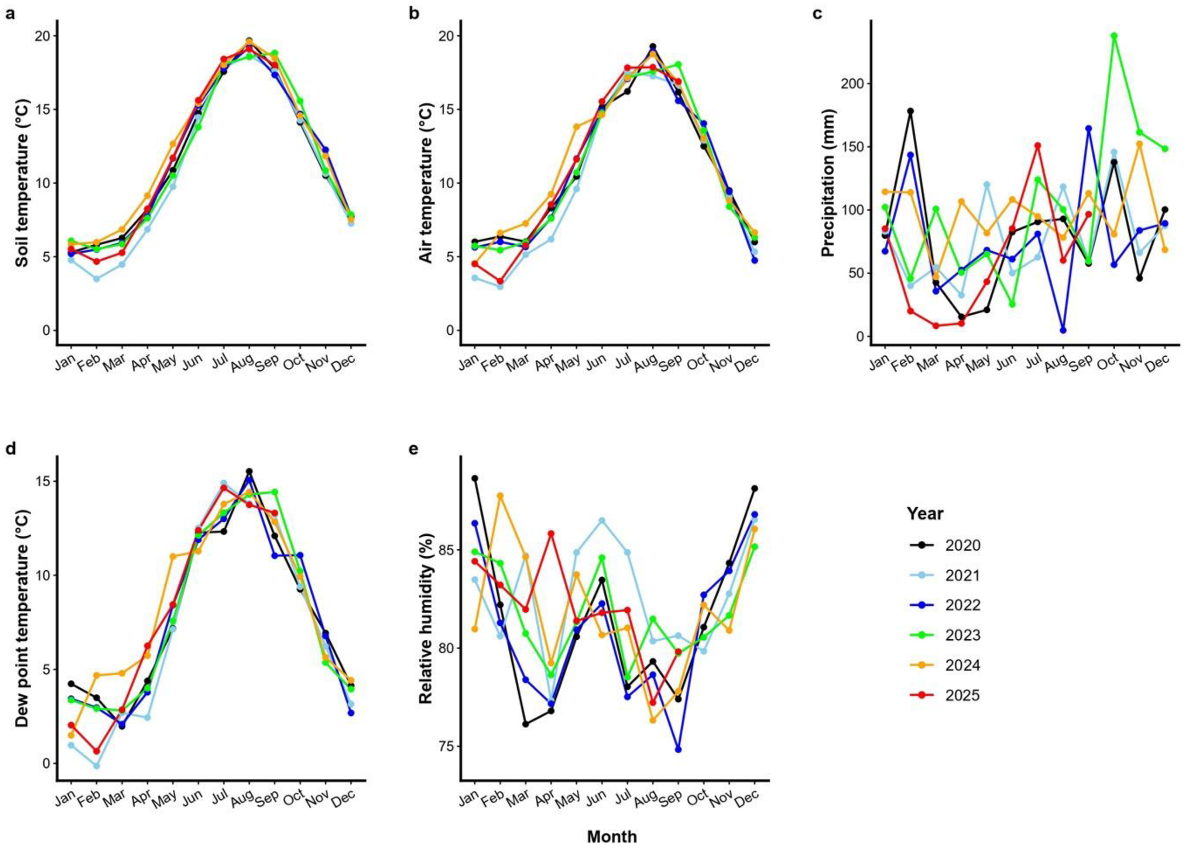


**Figure S13: Interannual variations in the mean monthly weather variables in Spiekeroog.**

Line plots of different climatic factors showing trends in monthly changes for each parameter at Spiekeroog (2020-2025). Displayed are (a) Soil, (b) Air, and (d) dew point temperatures, (c) precipitation, and (e) relative humidity trends.

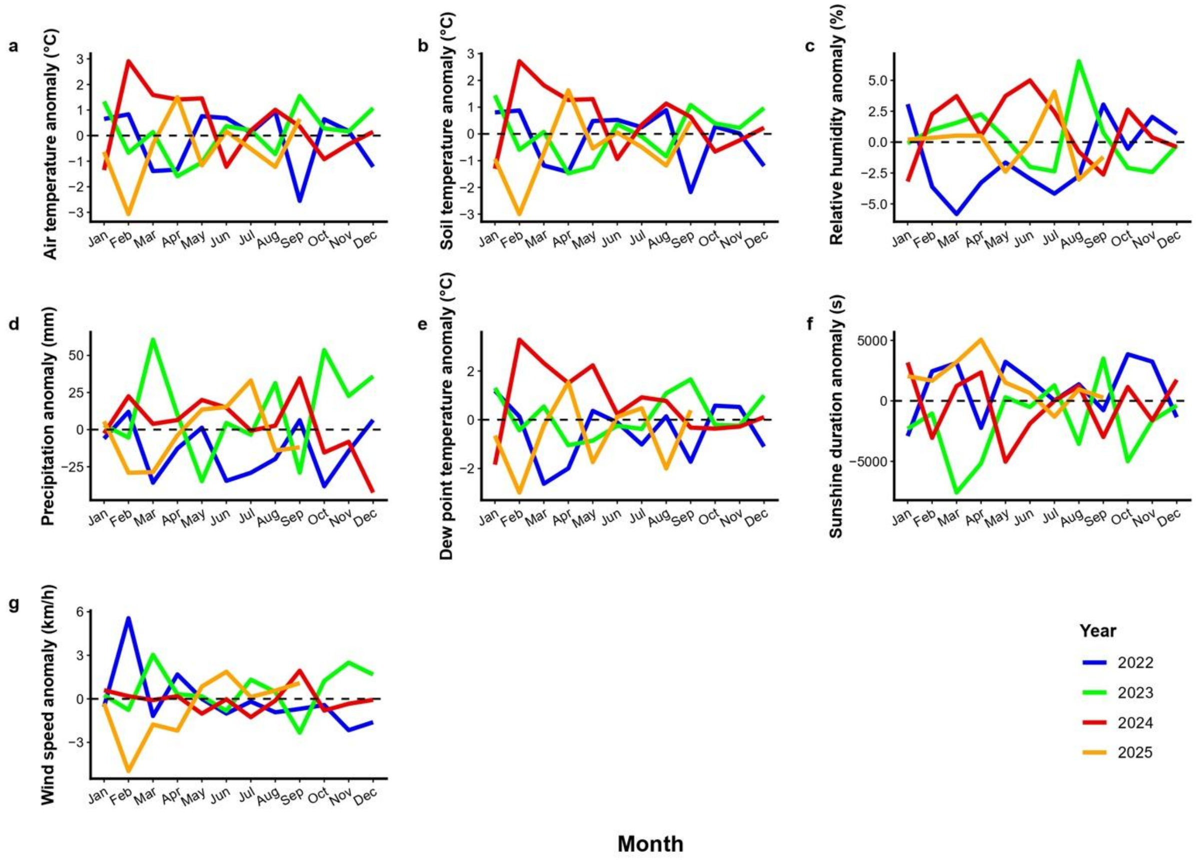


**Fig S14: Monthly weather anomalies at the Brachwitz collection site.**

Relative monthly anomalies in (a) air temperature, (b) soil temperature, (c) relative humidity, (d) precipitation, (e) dew point temperature, (f) sunshine duration, and (g) wind speed at the Brachwitz site across collection years (2022–2025). Anomalies were calculated as deviations from the corresponding monthly baseline means, enabling comparison of interannual variation in weather conditions across the sampling period.

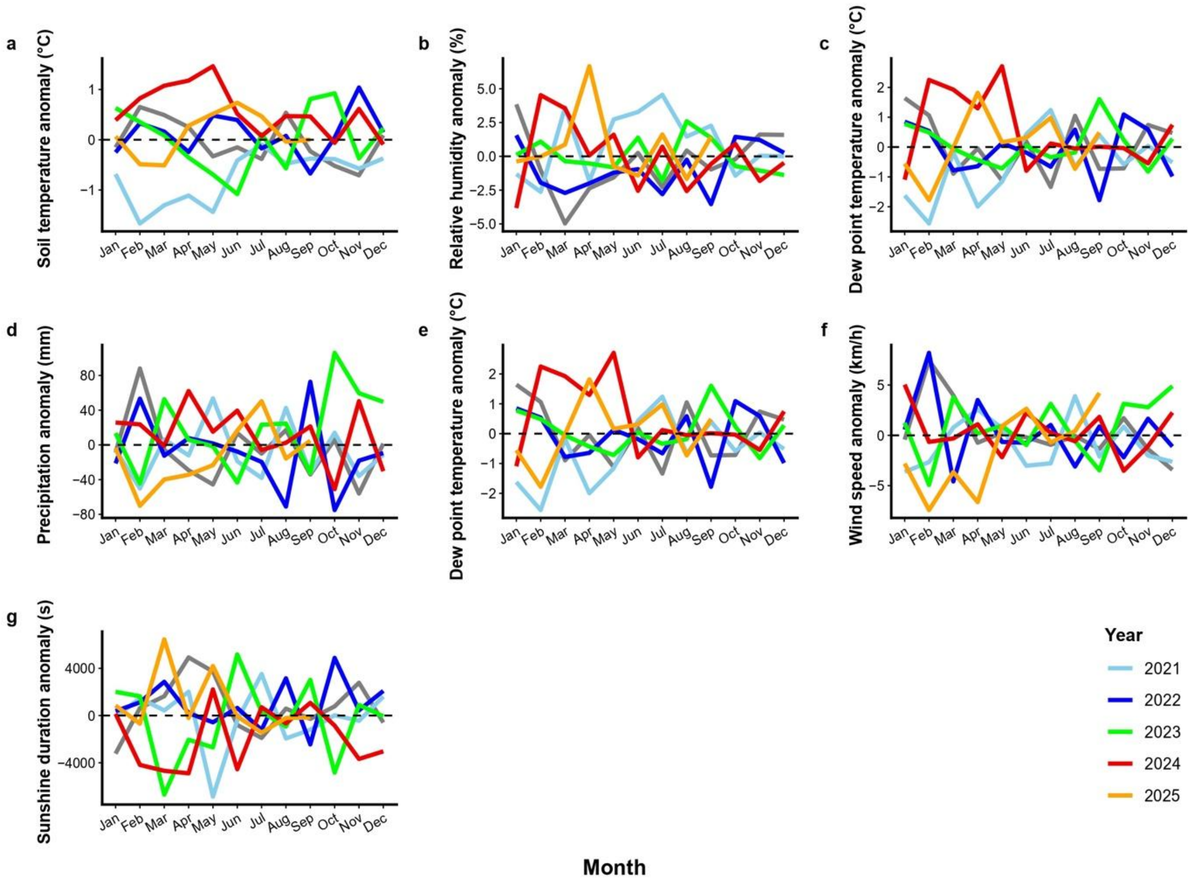


**Fig S15: Monthly weather anomalies at the Spiekeroog collection site.**

Relative monthly anomalies in (a) air temperature, (b) soil temperature, (c) relative humidity, (d) precipitation, (e) dew point temperature, (f) sunshine duration, and (g) wind speed at the Spiekeroog site across collection years (2021–2025). Anomalies were calculated as deviations from the corresponding monthly baseline means, enabling comparison of interannual variation in weather conditions across the sampling period.

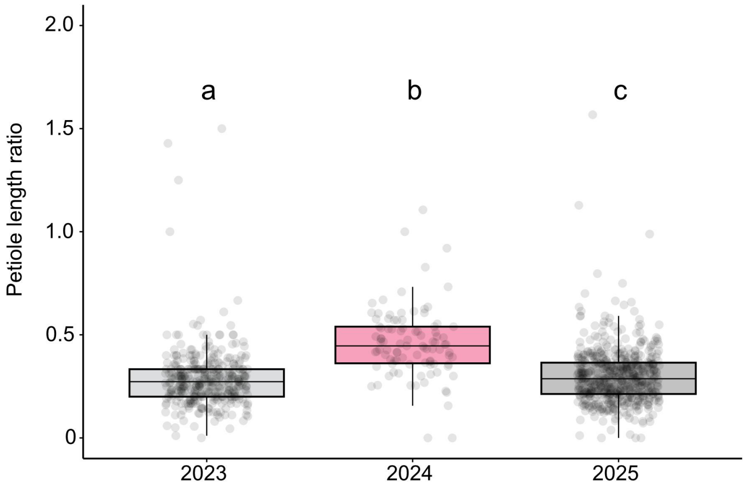


**Fig S16: Interannual temperature fluctuations influence the petiole elongation in wild A. thaliana.**

A box plot of the comparison in petiole length ratio of plants at Brachwitz across all collection years. The warmer year (2024) had plants with significant larger proportion of petiole length as compared to 2023 and 2025.

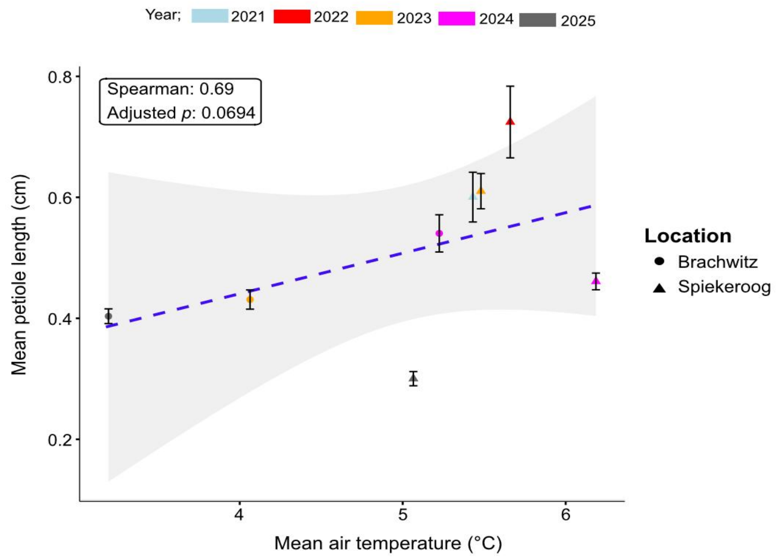


**Fig S17: Petiole length displays positive correlation with ambient air temperature.**

A line plot of mean petiole length for different years-locations collections (coloured points with error bars) against mean air temperature of the four months prior to April (sample collection). A regression line is fitted indicating the spearman's correlation between the petiole length and air temperature.

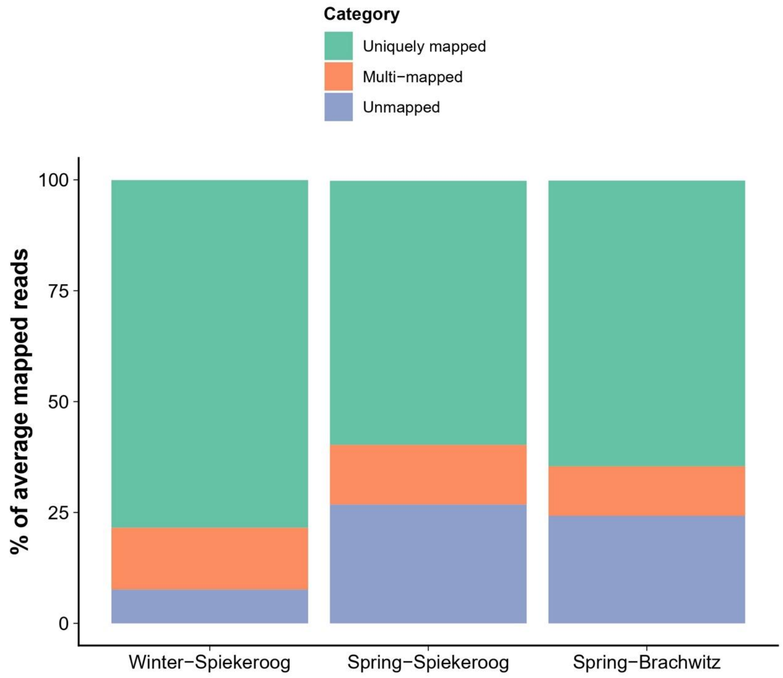


**Fig S18: Proportion of unmapped reads across seasons and location.**

A stacked bar plot displaying the percentage of the average mapped reads. Most reads aligned to the *A. thaliana* genome; however, a substantial fraction (10-30%) remained unmapped, with consistently higher proportions observed in spring than in winter samples.

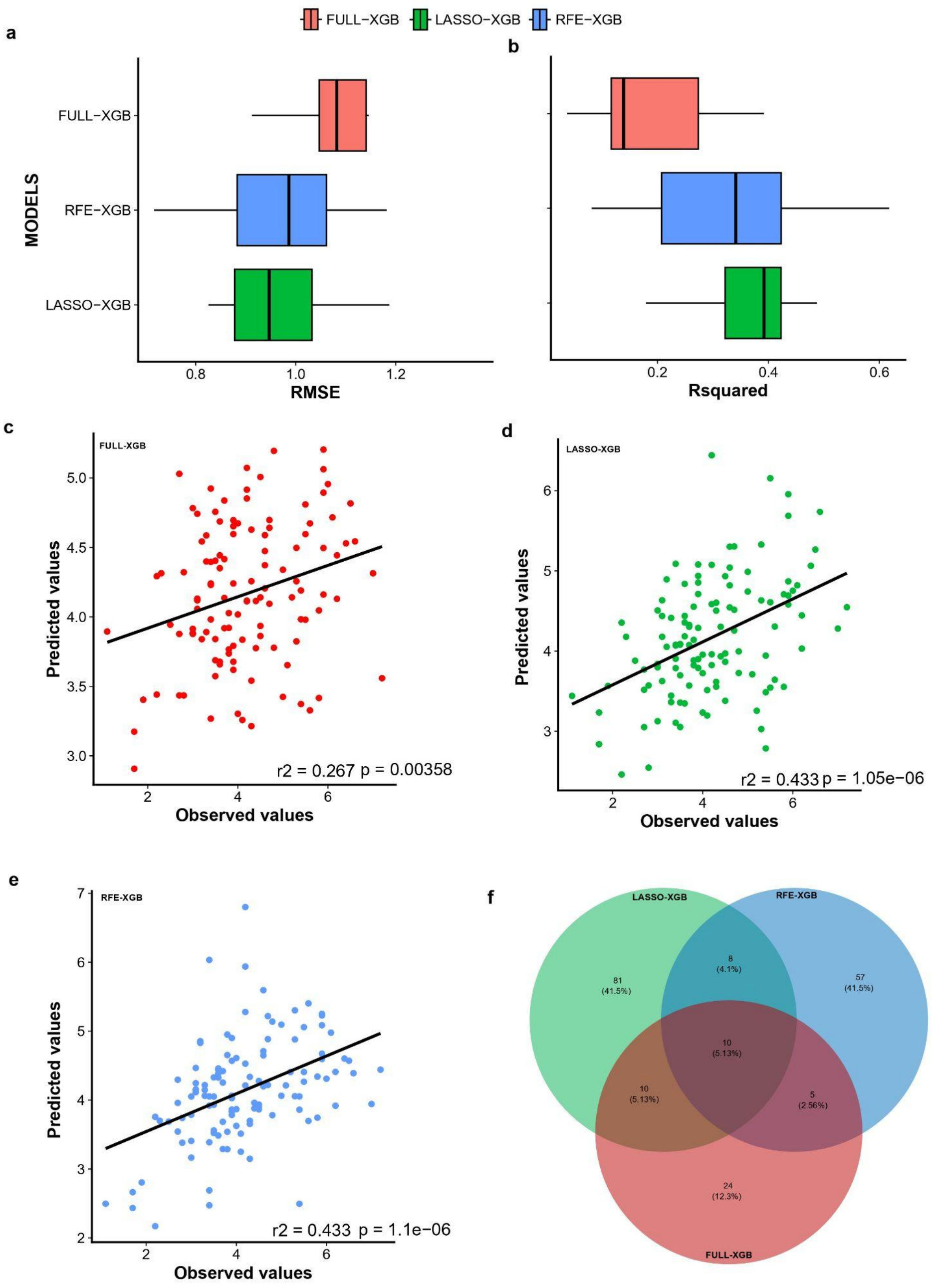


**Fig S19: Leaf surface temperature models comparative performance evaluation and gene overlap (Winter samples)**

**a-b:** Boxplots of cross-validated performance metrics (a) root mean squared error (RMSE) and (b) coefficient of determination ($R^2$) across 10 folds on the training and validation datasets. The boxes represent the interquartile range, with the median indicated by the central black line. The best-performing model (lower RMSE and higher $R^2$) is positioned toward the bottom of each panel. Associated with Data S15.

**c-e:** Point plots with fitted correlation lines for observed (from the testing dataset) versus predicted leaf surface temperature values from the FULL-XGB, LASSO-XGB, and RFE-XGB models. Pearson's correlation coefficient (r2) and the p-value for significance are indicated in each plot. Associated with Data S15.

**f:** A Venn diagram showing the proportion of unique important genes across the three models and their overlaps. Associated with Data S14.

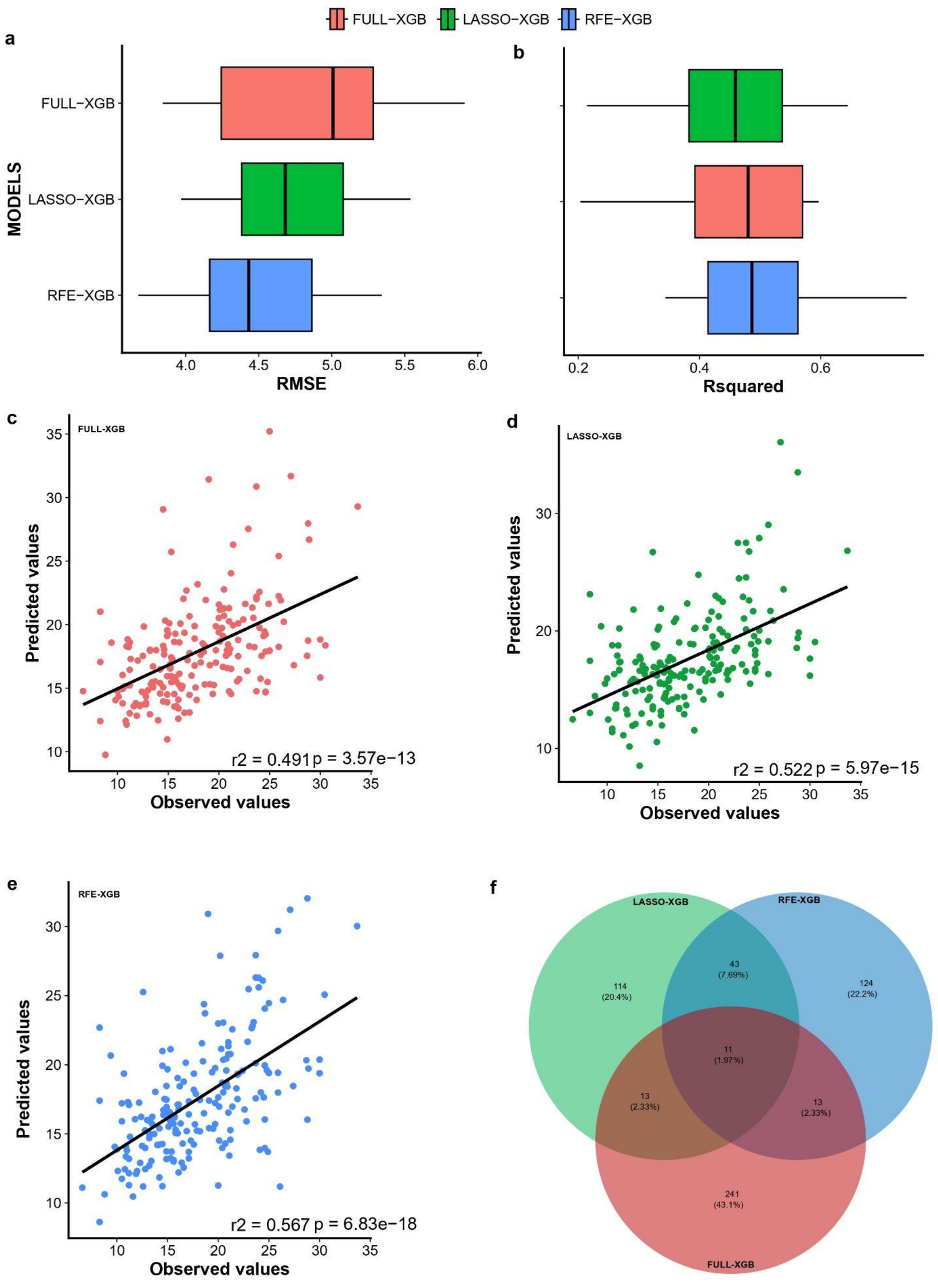


**Fig S20: Leaf surface temperature models comparative performance evaluation and gene overlap (Spring samples)**

**a-b:** Boxplots of cross-validated performance metrics (a) RMSE and (b) $R^2$ across 10 folds on the training and validation datasets. The boxes represent the interquartile range, with the median indicated by the central black line. The best-performing model (lower RMSE and higher $R^2$) is positioned toward the bottom of each panel.

**c-e:** Point plots with fitted correlation lines for observed (from the testing dataset) versus predicted leaf surface temperature values from the FULL-XGB, LASSO-XGB, and RFE-XGB models. Pearson's correlation coefficient (r2) and the p-value for significance are indicated in each plot. Associated with Data S16

**f:** A Venn diagram showing the proportion of unique important genes across the three models and their overlaps. Associated with Data S14.

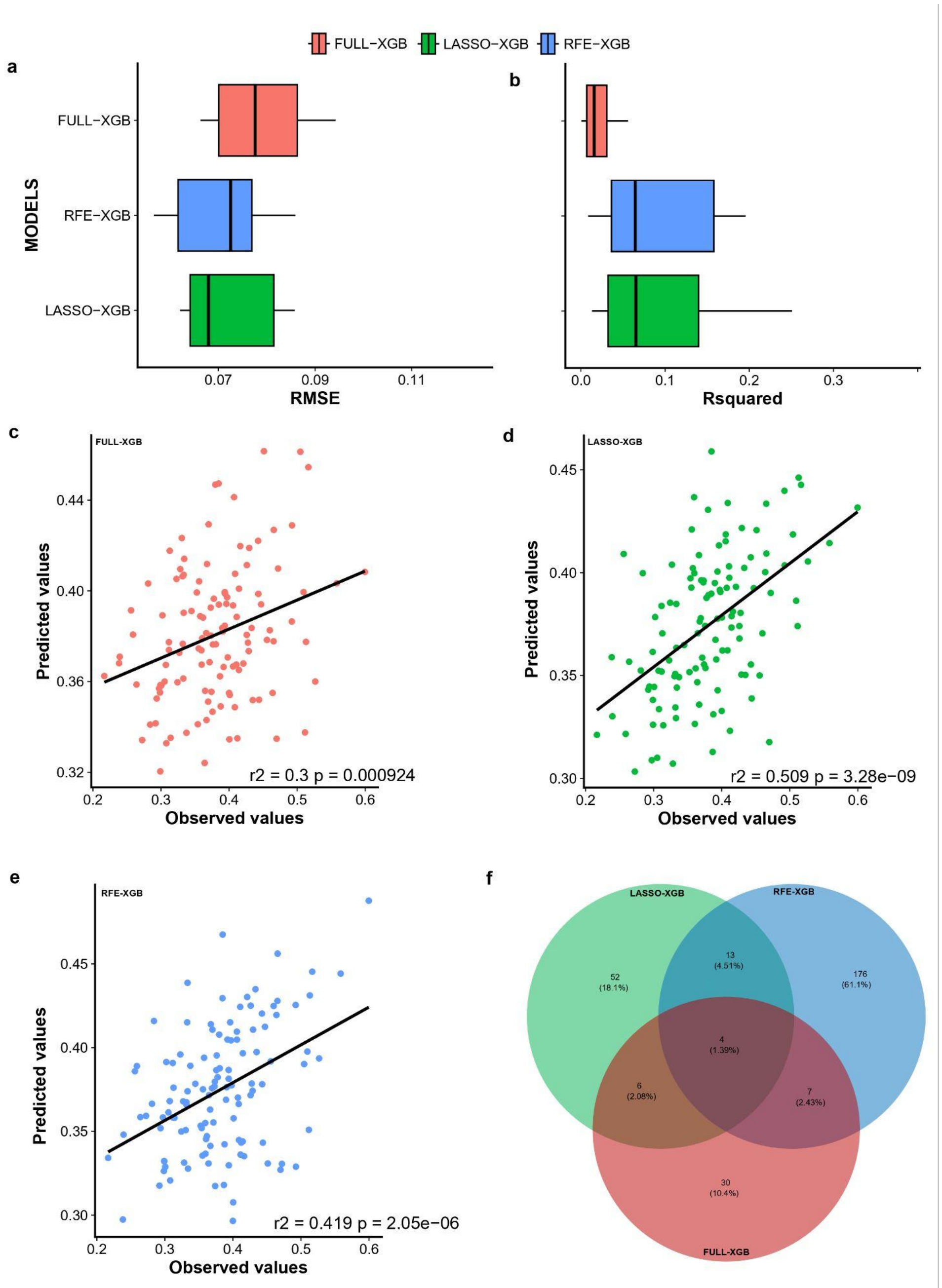


**Fig S21: Petiole length ratio models comparative performance evaluation and gene overlap (Winter samples)**

**a-b:** Boxplots of cross-validated performance metrics (a) RMSE and (b) $R^2$ across 10 folds on the training and validation datasets. The boxes represent the interquartile range, with the median indicated by the central black line. The best-performing model (lower RMSE and higher $R^2$) is positioned toward the bottom of each panel. Associated with Data S15.

**c-e:** Point plots with fitted correlation lines for observed (from the testing dataset) versus predicted petiole length ratio values from the FULL-XGB, LASSO-XGB, and RFE-XGB models. Pearson's correlation coefficient (r2) and the p-value for significance are indicated in each plot. Associated with Data S15.

**f:** A Venn diagram showing the proportion of unique important genes across the three models and their overlaps. Associated with Data S13.

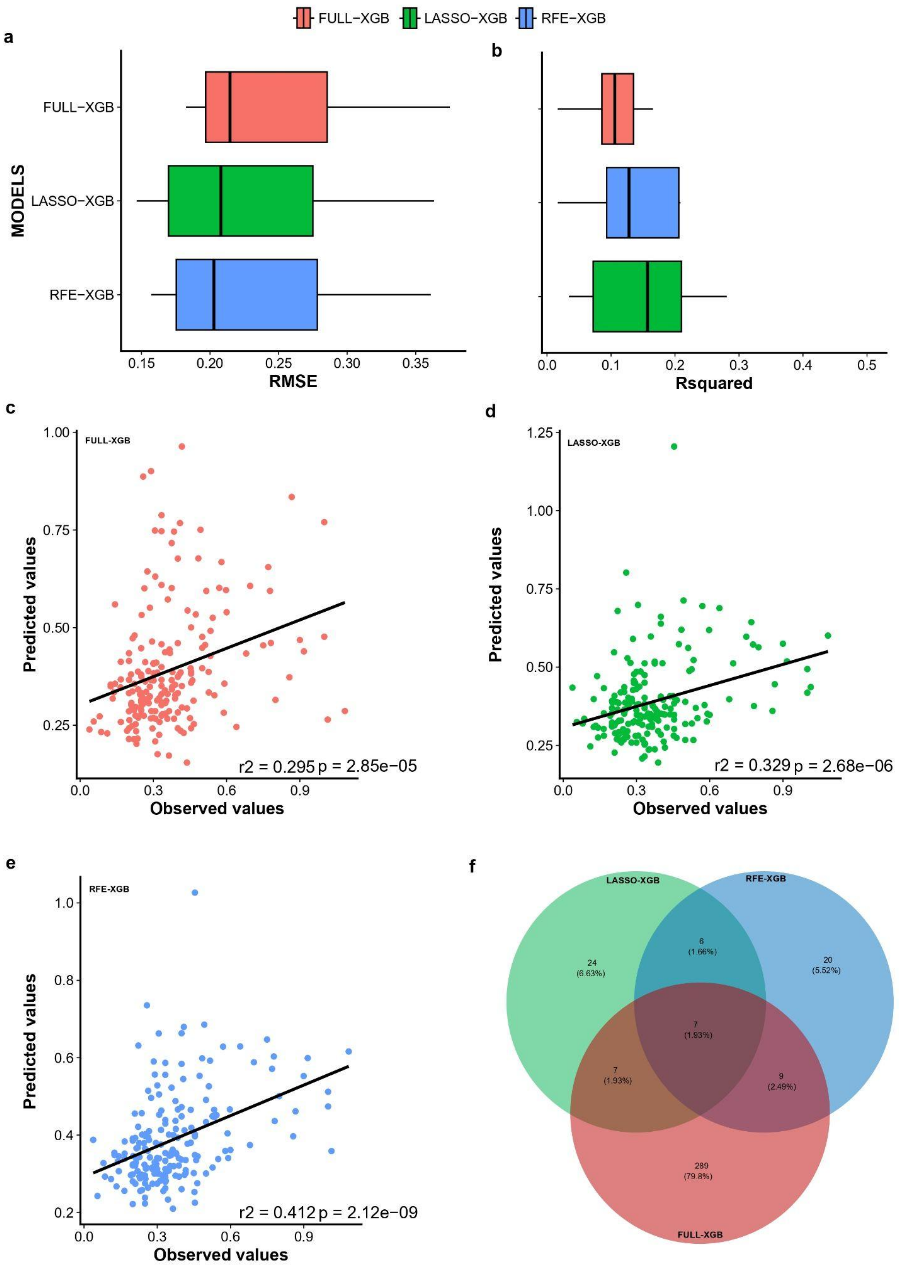


**Fig S22: Petiole length ratio models comparative performance evaluation and genes overlap (Spring samples)**

**a-b:** Boxplots of cross-validated performance metrics (a) RMSE and (b) $R^2$ across 10 folds on the training and validation datasets. The boxes represent the interquartile range, with the median indicated by the central black line. The best-performing model (lower RMSE and higher $R^2$) is positioned toward the bottom of each panel. Associated with Data S16.

**c-e:** Point plots with fitted correlation lines for observed (from the testing dataset) versus predicted petiole length ratio values from the FULL-XGB, LASSO-XGB, and RFE-XGB models. Pearson's correlation coefficient (r2) and the p-value for significance are indicated in each plot. Associated with Data S16.

**f:** A Venn diagram showing the proportion of unique important genes across the three models and their overlaps. Associated with Data S14.

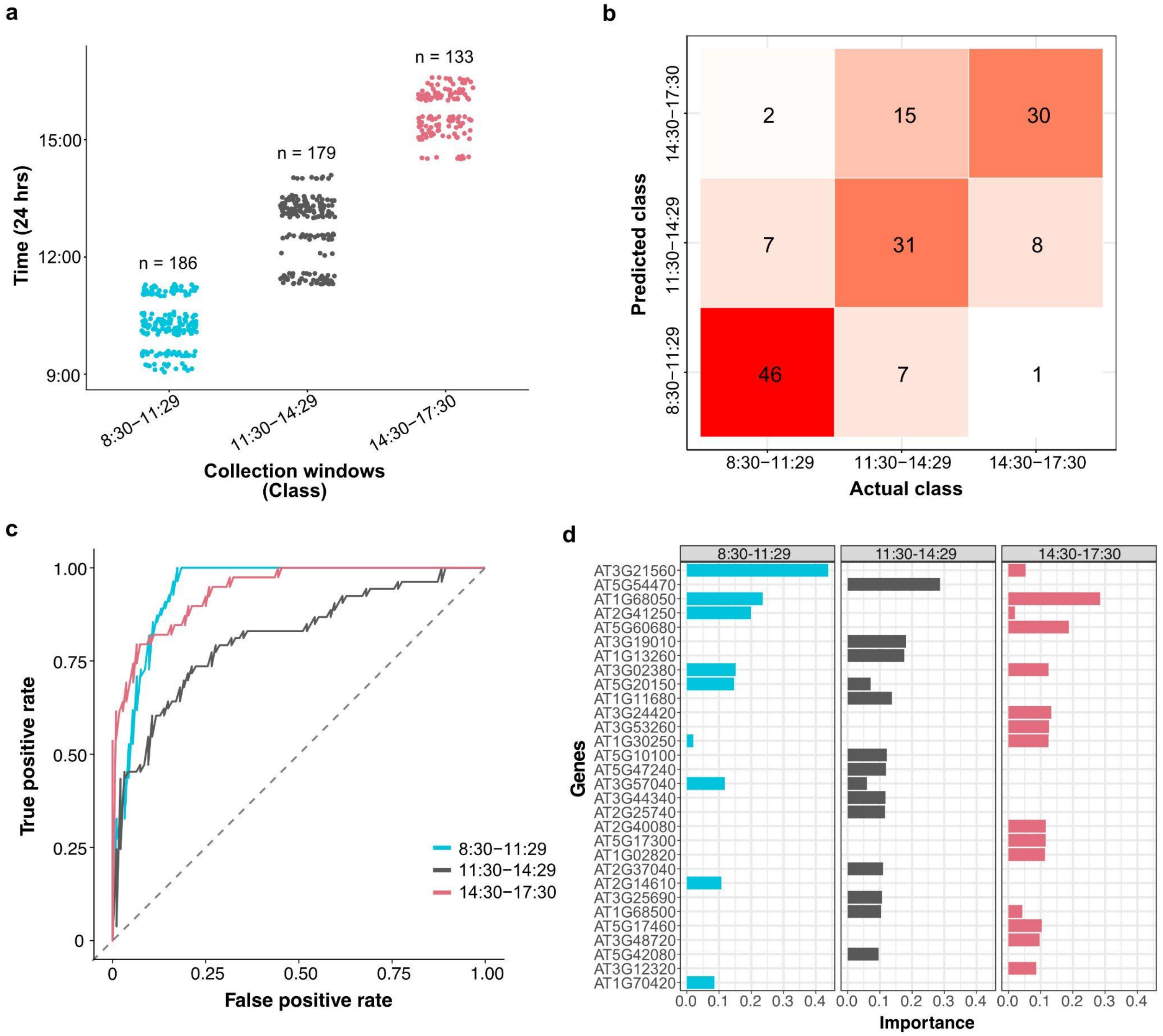


**Fig S23: Collection time associated expression profiles.**

**a:** Schematic overview showing the distribution of plant sample sizes collected in winter at Spiekeroog into their respective collection time bins. The time counts start from the sunrise (ZT0) for which on average was ~8:30 am on the collection days on the island.

**b:** A confusion matrix heatmap showing the number of predicted and actual classes for each time bin, highlighting the correct and misclassified instances. The classifier had better precision at predicting the first and last collection time as compared to the midday window (11:30-14:29). Specifically the model precision was 0.8032, 0.775 and 0.7173 for 8:30 am - 11:29 am, 11:30 am - 14:29 pm, 14:30 pm - 17:29 pm respectively.

**c:** Performance evaluation as depicted by the Receiver Operating Characteristic (ROC) curves indicating the true positive rate versus false Positive Rate. The true positive rate corresponds to the precision provided for each bin in panel b. The false positive rates were 0.9302, 0.7944 and 0.9406 for 8:30 am - 11:29 am, 11:30 am - 14:29 pm, 14:30 pm - 17:29 pm respectively. The lines are colored as indicated on the legend.

**d:** Top 30 important genes for predicting samples' collection bin ranked by their LASSO classification model coefficients. Notable genes which are under circadian regulation are AT2G40080 and AT1G68050. Associated with Data S16.

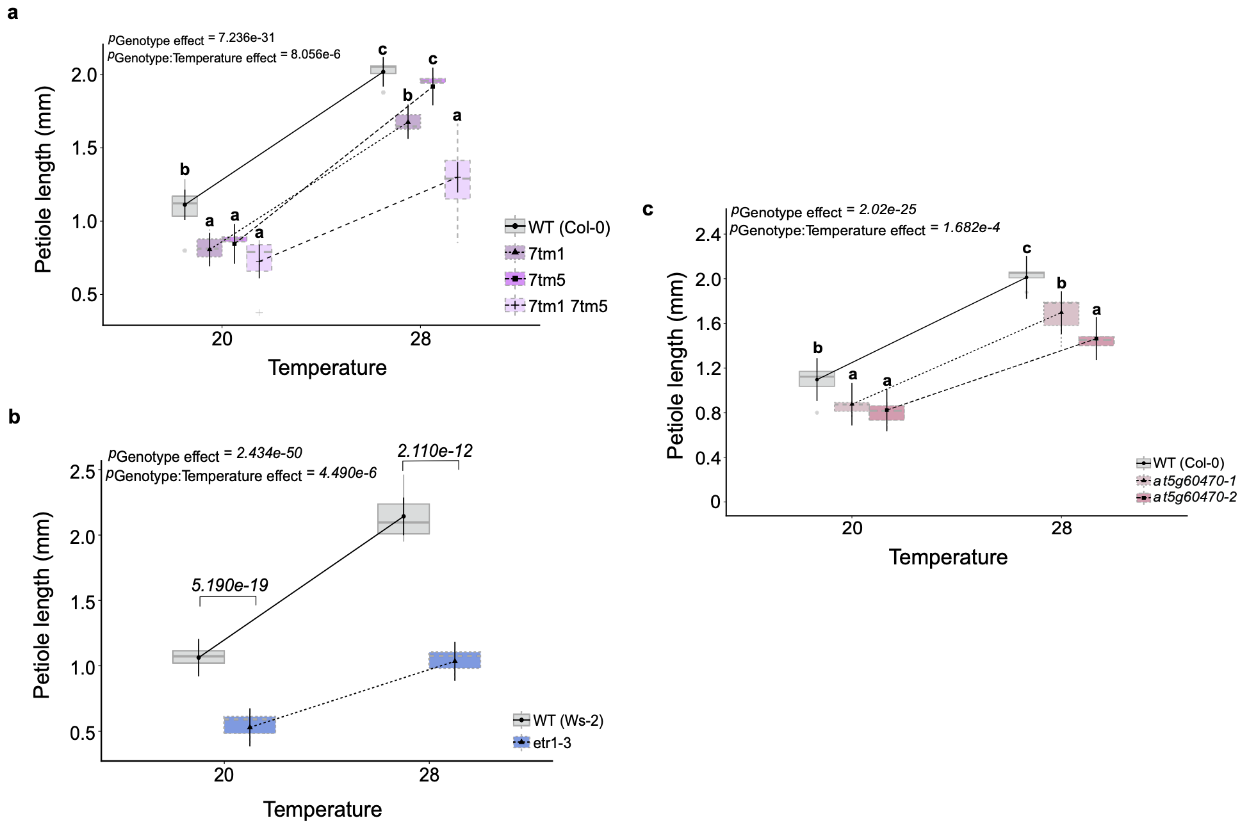


**Fig S24: Genotype-dependent influence on petiole growth and its interaction with temperature.**

**a-c:** Petiole length variation across multiple genotypes analyzed using linear mixed-effects models, with genotype and temperature included as fixed effects and experimental replicate modelled as a random effect. In these analyses, both the global genotype effect and the genotype × temperature interaction were significant and are provided on the top left on each plot, indicating genotype-specific temperature responses. Letters above boxplots denote post hoc groupings based on ANOVA, with shared letters indicating non-significant differences between genotypes within and across temperature conditions. These experiments are associated with Data S19.

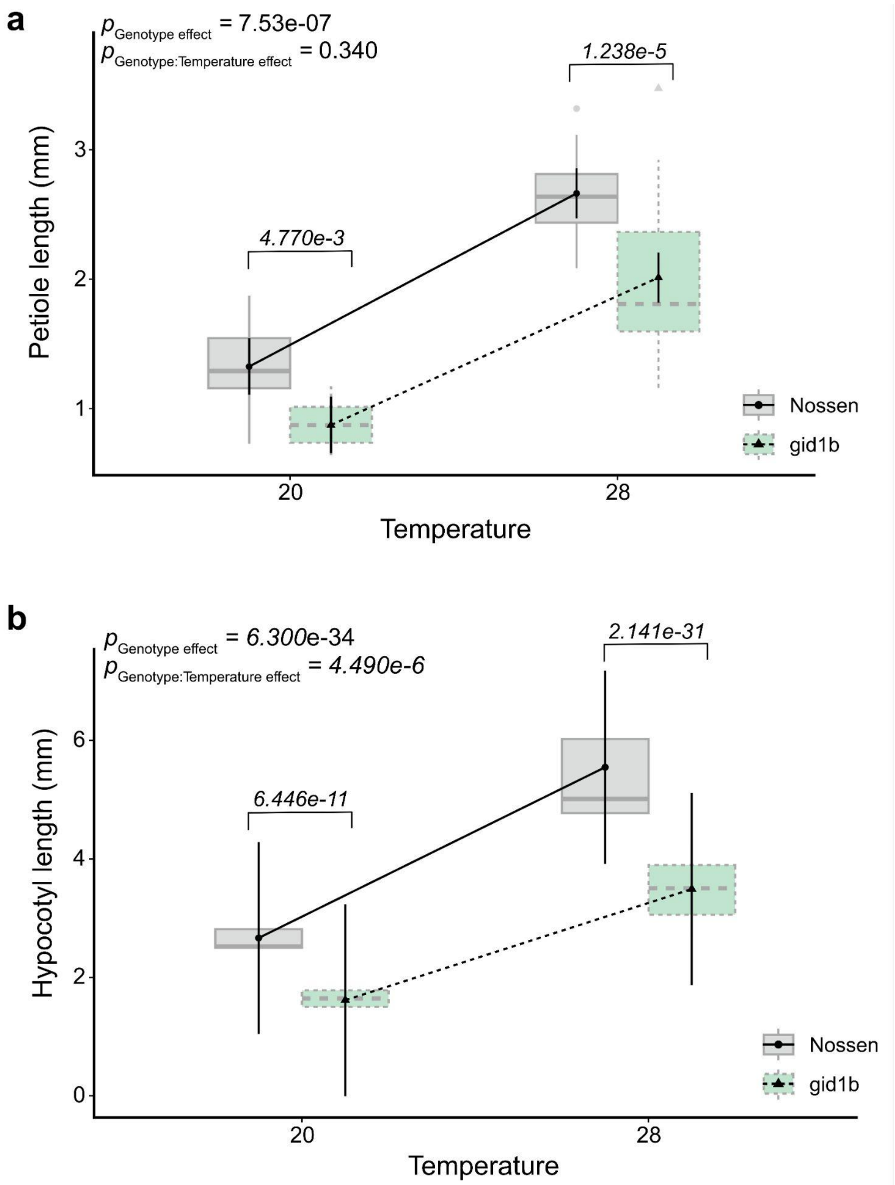


**Fig S25: *gid1b* mutants exhibited temperature-induced hypocotyl, but not petiole, elongation.**

Comparison between the WT (*Nossen*) and *gid1b* using a two-way ANOVA including genotype and temperature as fixed effects. Associated with Data S20.

**a:** A significant global genotype effect was detected in petiole elongation ($P_{\text{Genotype effect}} > 0.05$), whereas the genotype × temperature interaction was not significant.

**b:** Both genotype and genotype x temperature interaction showed significant effects on hypocotyl elongation.

**a. Plant ID**

Assign plant ID (tube label) - use a unique, pre-defined identifier

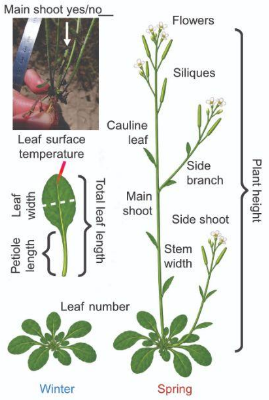


**b. Plant architecture**

☐Main shoot present: yes / no
☐Number of side shoots
☐Number of branches
☐Number of cauline leaves (total)
☐Plant height - measure from base to highest point of the main shoot (or main shoot)
☐Stem width - measure diameter of the main stem
☐Number of flowers (not fertilized; up to stage 14)
☐Number of siliques (fertilized flowers stage 15; elongated siliques)

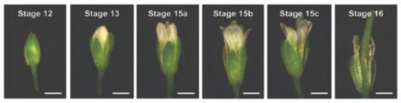


doi: 10.1104/pp.107.110908

**c. Rosette measurements**

☐Count number of rosette leaves - include all visible leaves
☐Identify longest rosette leaf - use this leaf for all subsequent steps
☐Leaf length - from base to tip
☐Leaf width - maximum width of the blade
☐Petiole length - from leaf base to blade

**d. Temperature (before sampling)**

☐Measure leaf surface temperature - use infrared thermometer on the longest leaf
☐Take a photo of the plant - include whole plant and surroundings
☐Ensure temperature reading is visible in the image
☐GPS coordinates and time will be extracted from image metadata
→ Check phone settings before taking the picture

**e. Sampling for RNA**

☐Remove longest rosette leaf (same leaf used for measurements), rinse briefly in water
☐Place immediately into labeled tube, gently pinch the leaf to allow RNAlater penetration, ensure full immersion in solution

**Fig S26: Field phenotyping and tissue collection protocol.**

**a:** A schematic representation of *A. thaliana* in winter and spring highlighting different traits measured during the field activities.
**b:** List of traits associated with plant architecture. These were mainly collected during spring season on the mature plants
**c:** Leaf-related traits. These were collected from both vegetative and reproductive/mature plants.
**d:** Leaf surface temperature measurements collected from the longest rosette leaf identified in c.
**e:** Leaf tissue collection for DNA/RNA isolation. The longest rosette leaf is detached from the sampled plant using a forcep.

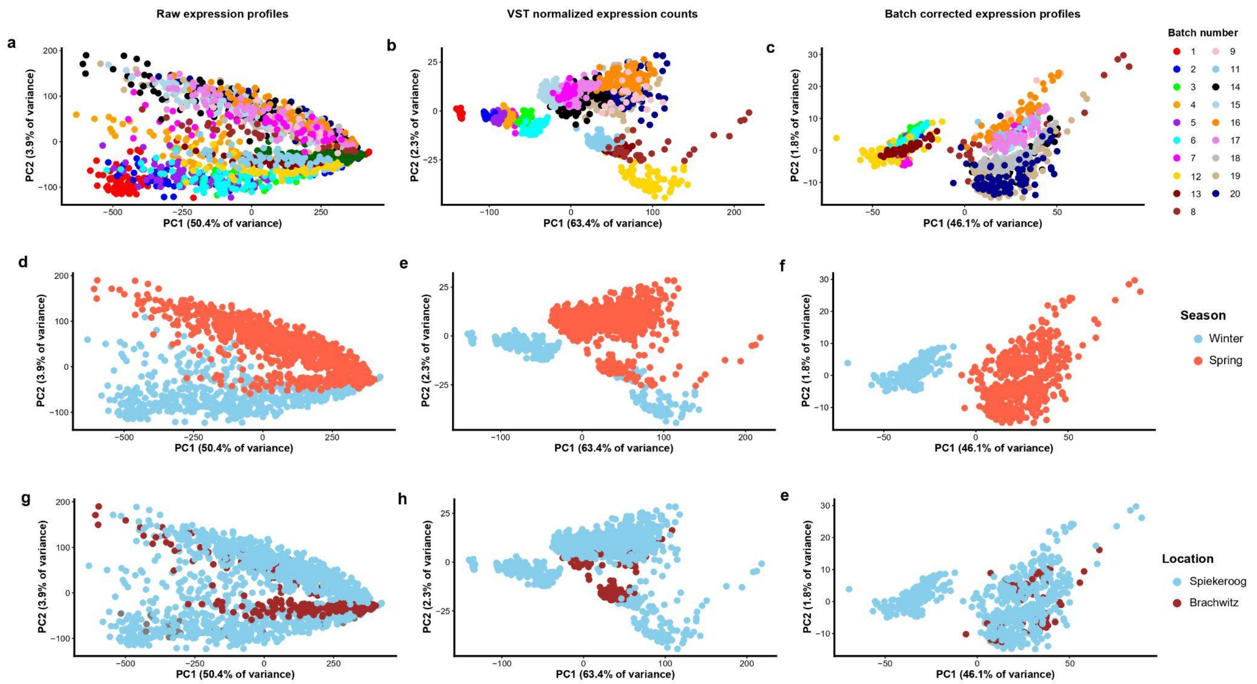


**Fig S27: Principal component analysis (PCA) of gene expression counts before and after normalization and batch correction.**

PCA biplot with the percentage of explained variation by each PC (PC1 and PC2) indicated on the axes. Plots (a, d, g), (b, e, h), and (c, f, i) correspond to raw, VST normalized, and batch corrected gene counts, respectively. Samples are coloured by batch number (a-c), season (d-f), and location (g-i). While variation associated mainly with season and location is preserved, the strong separation by batch observed in raw and normalized counts is largely removed after batch correction

#### Supplementary Data

**Data S1:** Description of the in situ phenotypic traits measured and metadata information recorded.

**Data S2:** All phenotypic data collected from 2021 to 2025 from different locations, mainly Brachwitz and Spiekeroog.

**Data S3:** Computed monthly means from hourly weather measurements for Brachwitz and Spiekeroog. Metadata for Supplementary Fig. S12 and S13.

**Data S4:** Calculated weather anomalies and metadata supporting Figure 2A and Supplementary Figs. 14 and 15.

**Data S5:** Weather variables and traits LMM results, highlighting significant weather-trait interaction and effect size. Metadata associated with Fig. 2c.

**Data S6:** Significant weather variable contributions to trait variation in our collection, as per RDA analysis. Metadata associated with Fig. 2d-e.

**Data S7:** Significantly up- and down-regulated genes from seasonal DGEs analysis. Metadata for Fig. 3d.

**Data S8:** Seasonal GO terms for biological processes. Metadata for Fig. 3d.

**Data S9:** Significant DEGs and GO terms for the locational DGE analysis. Metadata for Fig. 3f.

**Data S10:** Different trait measurements and genes used in prediction models for Winter samples

**Data S11:** Different trait measurements and genes used in prediction models for Spring samples

**Data S12:** Candidate genes for different traits from XGBoost models trained on Winter samples and metadata for Fig. 4b-d.

**Data S13:** Candidate genes for different traits from XGBoost models trained on Spring samples and metadata for Fig. 4b- d.

**Data S14:** Metadata for supplementary Figures 19 and 21. Performance metrics (RMSE, R2, and MAE) for all traits in winter.

**Data S15:** Metadata for supplementary Figures 20 and 22. Performance metrics (RMSE, R2, and MAE) for all traits in spring.

**Data S16:** Candidate genes corresponding to circadian rhythms.

**Data S17:** Metadata associated with petiole length validation on Fig 4f-h.

**Data S18:** Metadata associated with petiole length validation on Fig 4f-h

**Data S19:** Metadata associated with validation on the *gid1b* mutant in Fig. S24.

**Data S20:** Plant samples for DNA sequencing and metadata for Fig. 1b.

**Data S21:** BRB-seq library plates, their associated samples, and batches.

**Data S22:** Selected gene candidates from LASSO and RFE models for Winter

**Data S23:** Selected gene candidates from LASSO and RFE models for Spring
